# Driver-associated transcriptional rewiring reveals conditional genetic vulnerabilities in cancer

**DOI:** 10.1101/2024.11.20.624509

**Authors:** Sara Geraghty, Jacob A. Boyer, Mahya Fazel-Zarandi, Nibal Arzouni, Rolf-Peter Ryseck, Matthew J. McBride, Lance R. Parsons, Joshua D. Rabinowitz, Mona Singh

## Abstract

Mutations within cancer driver genes induce widespread transcriptional changes that reflect altered cellular states and can reveal associated genetic vulnerabilities. However, it remains challenging to determine which genes are dysregulated as a consequence of cancer alterations, and of these, which represent therapeutic opportunities. Here, we present Dyscovr, an integrative computational framework that leverages somatic mutation, gene expression, copy number alteration, methylation, and clinical data from primary tumors to identify driver-associated transcriptional changes. Dyscovr then uses these transcriptional changes as a biologically grounded starting point, integrating them with cancer cell line data to prioritize genes whose inhibition is predicted to reduce viability either specifically in driver-mutant contexts or in combination with driver inhibition. Applied both pan-cancer and across 19 tumor types, Dyscovr uncovers hundreds of such conditional vulnerabilities. As a case study, we newly implicate—and experimentally validate—*KBTBD2* as a gene whose inhibition enhances the efficacy of PI3K inhibitors in *PIK3CA*-mutant breast cancer cell lines. The Dyscovr software (github.com/Singh-Lab/Dyscovr) and predictions (dyscovr.princeton.edu) provide a platform and resource for linking mutated driver genes to conditional genetic vulnerabilities and for prioritizing these relationships for experimental and therapeutic investigation.

## INTRODUCTION

The personalized medicine approach to cancer treatment has largely focused on targeting an individual’s altered cancer driver genes. This strategy has shown considerable promise and is now routine in the clinic, enabled by the widespread use of mutational profiling panels (e.g. MSK-IMPACT^1^) and the emergence of drugs that specifically inhibit mutated driver genes^2^. However, not all patients with a targetable mutation respond to the corresponding therapy, and among those that do, tumors frequently acquire resistance, often through reactivation of the same oncogenic pathway. These limitations highlight the need for additional strategies to enhance the durability and effectiveness of personalized cancer treatments.

Mutations within driver genes affect more than just the gene itself, initiating cascades of molecular events that rewire transcriptional programs across the genome. These changes can reveal dysregulated genes that are functionally relevant to the driver-mutant context and that may also constitute candidate therapeutic targets. Because many genes may be dysregulated downstream of a driver alteration, one promising direction is to identify which of these genes participate in genetic interactions with the driver, particularly those that represent conditional genetic vulnerabilities such that their inhibition compromises cell viability either in the context of a somatically altered driver or in combination with pharmacologic inhibition of the driver. Focusing on transcriptionally dysregulated genes is advantageous: because transcriptional rewiring reflects the functional state of tumor cells, it provides a biologically grounded starting point for discovering such conditional genetic vulnerabilities, rather than considering all possible genes. Accordingly, a systematic computational approach that first uncovers driver-associated transcriptional changes and then evaluates the dysregulated genes for genetic interactions offers a principled path for identifying both individual drug targets and co-targets that may enable more durable therapeutic responses.

Here, we introduce Dyscovr, an approach for identifying genes that are transcriptionally dysregulated and that participate in driver-associated genetic interactions. Dyscovr first identifies robust driver-associated transcriptional changes by integrating multiomic data from primary tumors, while accounting for molecular and clinical covariates that influence baseline gene expression. It then leverages functional genetic screening data from cancer cell lines to determine which dysregulated genes genetically interact with the driver and to prioritize the subset that represents candidate conditional genetic vulnerabilities with therapeutic potential.

Despite major progress in cancer systems biology, to our knowledge, no existing approach jointly links driver mutations to their genome-wide transcriptional consequences in a way that enables the discovery of genetic interactions with the driver. Prior research has instead examined related problems within these two areas in isolation, leaving the relationship between driver-associated transcriptional rewiring and conditional genetic vulnerabilities largely unexplored. A significant portion of research relating somatic mutations to expression changes has focused on identifying cancer genes or pathways by assessing whether direct interaction partners or network-proximal genes are dysregulated^3–10^. These approaches typically rely on *a priori* knowledge of regulatory or protein-protein interaction networks, which can restrict analyses to predefined relationships and limit their ability to capture the impact of gene-level somatic alterations on the full transcriptome. Other approaches model the impact of somatic mutations on regulatory activity, such as transcription factor activity^11,12^, providing insight into altered regulatory programs but without directly relating somatic mutations to quantitative expression changes across all genes. Simpler statistical tests have also been proposed to identify genes with altered expression with respect to somatic mutations in other genes^13^, but do not account for key confounding factors that affect gene expression in cancer, such as copy number alteration (CNA) or DNA methylation changes. Finally, while expression quantitative loci (eQTL) analysis─a cornerstone of gene regulation and variant impact research that has shown great success in linking individual variants to a gene’s expression levels^14^─has also been applied in the cancer context^15^, it is primarily designed to link germline variants to gene expression changes and can be difficult to apply to somatic mutations, which are rarer, heterogeneous across tumors, and often confounded by other genomic alterations. As a result, previous approaches do not provide a complete view of the global transcriptional impact of somatic mutations in cancer or systematically nominate dysregulated genes for further therapeutic investigation.

A separate line of research has aimed to uncover genetic vulnerabilities in cancer. This work has largely focused on identifying synthetic lethal (SL) interactions, in which inactivation of one gene (e.g. a mutated tumor suppressor) renders the other essential, and synthetic dosage lethal interactions, where overexpression of one gene (e.g. an oncogene) renders the other gene essential^16^. Approaches for identifying these interactions include supervised learning on known SL interactions^17–19^, statistically uncovering signals such as mutual exclusivity of genetic alterations in human tumor data^20–22^, and analyzing dependencies in cell line screens^23–25^. However, these methods do not link predicted vulnerabilities to the transcriptional consequences of driver gene mutation, and many are not grounded in molecular data from primary patient tumors. Further, in contrast to previous approaches, Dyscovr distinguishes between two classes of driver-associated genetic interactions with distinct therapeutic implications: (i) cases in which a mutated driver gene induces or enhances a dependency on a transcriptionally dysregulated gene, nominating that gene as a target for a single-agent strategy in that molecular context (i.e. “acquired vulnerabilities”); and (ii) cases in which joint inhibition of the driver and the dysregulated gene negatively impacts cell growth, indicating a co-targeting strategy (i.e. “co-targeting vulnerabilities”). In particular, the second class nominates candidate combination strategies involving joint inhibition of an oncogenic driver and a transcriptionally dysregulated partner gene to more effectively suppress tumor growth. Because many driver alterations involve activating mutations in oncogenes rather than loss of tumor suppressors, approaches that can identify such co-targeting relationships are critical, yet these relationships are not directly captured by traditional cancer SL discovery methods.

Dyscovr is a modular framework composed of two main stages. In the first stage, Dyscovr leverages multiomic data from The Cancer Genome Atlas (TCGA) and uses an integrative linear regression framework to uncover genes whose expression is significantly altered in tumors with mutations in a given driver gene. Conceptually, Dyscovr builds on eQTL analysis in several key ways: it models gene-gene associations by aggregating nonsynonymous somatic mutations across sites within a gene, accounts for the effects of multiple mutated genes within the same model, and integrates numerous other molecular and clinical factors that are critical for the cancer context. In the second stage, Dyscovr uses genome-wide knockout screening data from the Cancer Dependency Map (DepMap)^26^ to determine which of these transcriptionally altered genes exhibit genetic interactions with the driver gene and, among these interactions, which correspond to conditional genetic vulnerabilities with therapeutic potential. Importantly, once these relationships are learned from TCGA and DepMap data, they can be used to prioritize candidate therapeutic vulnerabilities based solely on driver mutation profiles, which are routinely obtained in clinical settings.

Across 19 cancer types, Dyscovr reveals hundreds of significant and novel associations between driver genes and transcriptionally dysregulated targets, including a subset predicted to participate in genetic interactions with the driver gene and to represent previously uncharacterized candidate therapeutic vulnerabilities. We demonstrate the power of Dyscovr in breast cancer by identifying and validating a genetic interaction between *PIK3CA* and *KBTBD2,* a gene we implicate as a cancer-relevant positive regulator of the PI3K-AKT pathway and insulin signaling. Our experiments show that inhibiting *KBTBD2* enhances the efficacy of PI3K inhibitors specifically in *PIK3CA*-mutant breast cancer cell lines, suggesting a new combinatorial strategy for treating *PIK3CA*-mutant breast tumors.

Overall, Dyscovr provides a systematic framework for connecting mutated cancer driver genes to downstream transcriptional dysregulation and for prioritizing potential driver-specific therapeutic vulnerabilities, including both candidate single-agent targets and co-targets for combination therapy. By integrating tumor multiomic data with functional dependency screens, Dyscovr identifies genetic interactions linked to specific oncogenic alterations, offering a principled approach for nominating combinatorial therapeutic strategies. Dyscovr is available both open source (github.com/Singh-Lab/Dyscovr) and as an interactive web resource (dyscovr.princeton.edu), facilitating future experimental efforts to translate these findings into therapeutic strategies.

## RESULTS

### Overview of Dyscovr Framework

Dyscovr is a framework that both links somatically mutated cancer driver genes to transcriptional changes and subsequently identifies conditional genetic vulnerabilities (Fig. 1). Dyscovr integrates matched mutation, CNA, methylation, and expression data from primary tumor samples from the TCGA (Fig. 1A) and focuses on the set *S* of known cancer driver genes^27^ that frequently (≥5%) possess nonsynonymous mutations in the given patient cohort (Fig. 1B). This restriction ensures that each driver has sufficient mutated samples for reliable association testing while focusing the analysis on mutations most likely to alter protein function.

**Figure 1.**
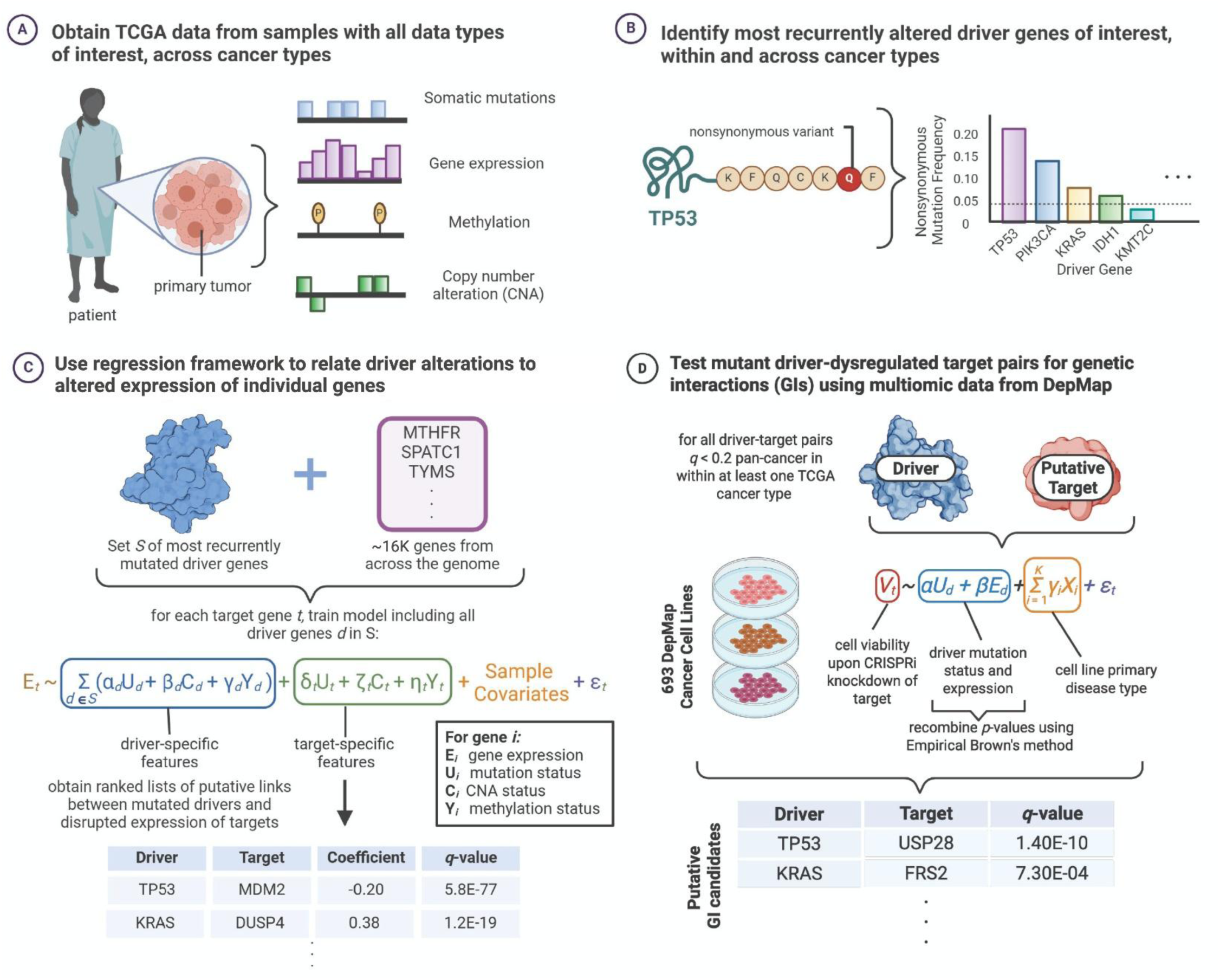
Methodological overview of Dyscovr pipeline. **A.** Dyscovr leverages matched somatic mutation, gene expression, methylation, and copy number alteration (CNA) data from primary patient tumors, such as those available through the Cancer Genome Atlas (TCGA). **B.** Across all cancer types, and within individual cancer types, Dyscovr considers a set *S* of the most recurrently mutated cancer driver genes across patient samples. Nonsynonymous mutations, including missense, nonsense, nonstop, and splice site mutations, are considered. **C.** Within each sample population, Dyscovr fits a linear regression model for each candidate target gene *t* and tests for an association between the expression of *t* and the nonsynonymous mutation status of driver genes. The model also incorporates other driver– and target-specific features, including CNA and methylation status, as well as clinical features. The expression of gene *t* is correlated to the nonsynonymous mutation status of a driver gene *d* if the fit coefficient α_*d*_ is significantly different from zero (Methods I). **D.** For cases where Dyscovr identifies a significant association between the nonsynonymous mutation status of a driver gene *d* and the expression of a target gene *t*, both in a pan-cancer context and within at least one individual cancer type, we evaluate the pair for a potential genetic interaction using cell line data from the Cancer Dependency Map (DepMap)^26^. We use a linear regression model to test for an association between cell viability upon CRISPR knockdown of target gene *t* and both the nonsynonymous mutation status and expression of *d* (Methods VII). This process results in a ranked list of target genes that Dyscovr predicts to participate in a genetic interaction with the corresponding driver gene; these interactions nominate driver-associated conditional genetic vulnerabilities.

In the first phase, for each candidate gene in the human genome, Dyscovr fits a linear regression model that jointly estimates the association between the gene’s expression and the nonsynonymous mutation status of each driver gene *d* in *S*. The model also accounts for other factors that may influence gene expression, including: CNA and methylation status of each driver; the putative downstream gene’s mutation, CNA, and methylation status; and sample-level covariates (e.g. patient age, sex, treatment status, tumor subtype, estimated fraction of infiltrating immune cells, genotypic variation, etc.) (Fig. 1C, Methods I-IV). For each regression performed, Dyscovr extracts the fit coefficients corresponding to driver mutation status and performs per-driver multiple hypothesis testing correction, yielding a ranked set of associations between mutated driver and dysregulated target genes with associated *q*-values, estimated magnitudes, and directionalities. We apply this framework both pan-cancer and within each of the 19 TCGA cancer types with at least 75 samples possessing all required data types, using *q* < 0.2 as a significance threshold. Because *q*-values tend to be smaller with an increased number of tumor samples considered, we report results for the pan-cancer analysis using a *q* < 0.01 threshold.

In the second phase, Dyscovr leverages these predicted transcriptional dysregulation effects to uncover genes that are involved in genetic interactions with the associated cancer driver genes and, of these, which constitute conditional genetic vulnerabilities with therapeutic potential (Fig. 1D, Methods VIII). For each driver-target pair identified as significant in the first part of Dyscovr—both pan-cancer and within at least one individual cancer type—we fit a second linear regression model to evaluate whether driver gene activity (measured by nonsynonymous mutation status and gene expression levels) influences cell viability following knockout of the target gene, while accounting for cancer type. In the absence of a genetic interaction, driver activity is not expected to affect cell viability upon target gene knockout. We apply this model across 693 cancer cell lines from the Cancer Dependency Map (DepMap), using *q* < 0.2 as our significance threshold.

### Dyscovr’s Predicted Dysregulated Targets are Enriched in Cancer-Related Genes and Known Functional Interactors

We first apply the initial stage of the Dyscovr framework to uncover relationships between nonsynonymous mutations in recurrently mutated cancer driver genes and the expression of 16,447 putative target genes with sufficient data and variability across patient samples (Methods IV.A, Data S1). Although Dyscovr models multiple molecular features, we focus our interpretation on the mutation coefficients for each driver gene. This reflects both the routine clinical availability of mutational data in tumor profiling and our goal of generating mutation-centered predictions that are feasible to validate and incorporate into therapeutic strategies.

Across primary samples spanning 32 cancer types in the TCGA, four annotated cancer driver genes^27^ are mutated at greater than 5% frequency overall and in at least two cancer types: *TP53*, *PIK3CA*, *KRAS*, and *IDH1* (Table S1). For each of *TP53*, *PIK3CA*, *KRAS*, and *IDH1*, Dyscovr identified hundreds to thousands of downstream target genes whose expression is significantly correlated to each driver’s nonsynonymous mutation status (Fig. 2A). *TP53*, the most highly mutated driver gene across TCGA samples, accounts for the largest number of dysregulated target genes, or ‘hits’, at a *q* < 0.01 significance threshold. Our results suggest Dyscovr prioritizes each driver’s known targets or functional partners: for *TP53, PIK3CA, KRAS,* and *IDH1*, each gene’s hits were significantly enriched in either the driver’s transcriptional targets from DoRothEA^28^, if a TF (i.e. *TP53*), or in the transcriptional targets of downstream TFs that have been shown the literature to be intermediaries in enacting their effects (i.e. *PIK3CA, KRAS, and IDH1*) (Fig. 2B, Table S2). *TP53* also has exceptionally well-classified downstream targets from a variety of other sources, and we find that the hits Dyscovr identifies for *TP53* are significantly enriched for curated *TP53* targets^29^ (Fig. 2C) as well as for targets identified in the TF-target databases TRRUST^30^ and hTFtarget^31^ (Table S3). *TP53*’s hits are also significantly enriched in established pathway genes from KEGG (hsa04115)^32^ and Reactome (R-HSA-3700989, R-CFA-5628897) (Fig. 2C, Table S3). Taken together, these strong enrichments across a variety of sources suggest that Dyscovr is effectively capturing transcriptional changes in downstream genes, including both direct transcriptional targets and pathway targets that may lie further downstream of the mutational event. Notably, approximately 72% of Dyscovr’s 100 most significant *TP53*-associated hits are not described in any of these curated sources, suggesting that Dyscovr can also nominate previously uncharacterized regulatory relationships even for the most well-studied cancer driver genes.

**Figure 2.**
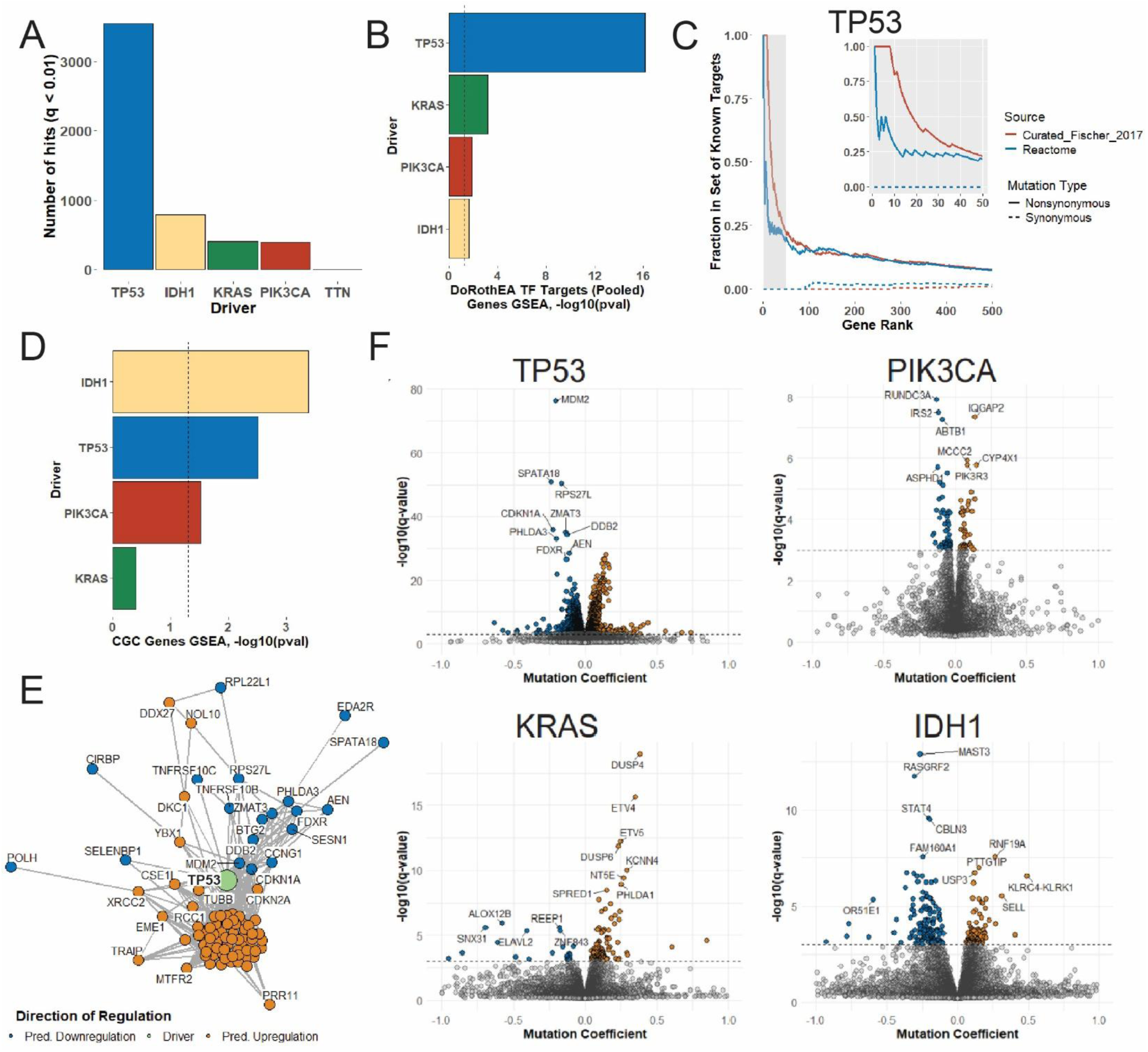
Dyscovr uncovers pan-cancer associations between mutated drivers and target gene expression changes. **A.** The number of significant associations (*q* < 0.01) identified for *TP53*, *PIK3CA*, *KRAS*, and *IDH1*, the four cancer driver genes included in pan-cancer modeling. Each hit represents a putative relationship between a nonsynonymous mutation in the driver gene and an expression change in a target gene. The first stage of Dyscovr uncovers hundreds of significant associations at this threshold for all recurrently mutated driver genes, while yielding only three hits for *TTN* (a frequently mutated gene not annotated as cancer-relevant^27,33^). **B.** Bar chart showing the enrichment of effector TF targets from DoRothEA^28^. For *TP53,* enrichment is calculated using *TP53-*specific target genes (N = 248). For *PIK3CA, KRAS*, and *IDH1,* enrichment is calculated using pooled targets from multiple literature-supported effector TFs (*PIK3CA:* N = 605; *KRAS:* N = 679; *IDH1:* N = 240; Table S2). Bar height represents *-log10*(*p*-value) from a one-sided GSEA test, with dotted line indicating *p* = 0.05. **C.** The cumulative fraction of curated *TP53* targets (Fischer et al.^29^, blue) and members of the *TP53* signaling pathway (Reactome^41^ R-HSA-3700989, red) among an increasing fraction of *q-*value-ranked *TP53* hits from a Dyscovr model using nonsynonymous mutations (solid lines) or silent mutations (dashed lines) as a control. Inlay (top right) shows the top 50 ranked *TP53* hits. **D.** Bar chart showing the enrichment of known cancer genes from the Cancer Gene Census (CGC)^33^ among each driver gene’s *q*-value ranked targets from Dyscovr. Bar height represents *-log10*(*p*-value) from a one-sided GSEA test, with the dotted line indicating *p* = 0.05. **E.** Network visualization of interconnected Dyscovr stage one results for *TP53* overlaid on the STRING functional protein-protein interaction network^35^, with *TP53* shown in green. The top 100 *TP53* hits (*q* < 0.01) connected either to *TP53* or to another hit with STRING confidence >0.4 are displayed. Node color indicates the predicted direction of regulation (upregulation in orange, downregulation in blue). **F**. Volcano plots of Dyscovr predictions for each driver gene, with a selection of the most statistically significant hits labeled. The *x*-axis shows the regression coefficient for the driver mutation status term (thresholded to the range [-1, 1] for visualization), with predicted upregulated target genes in orange and downregulated target genes in blue at *q* < 0.01 (dotted line). The *y*-axis shows the *-log10*(*q*-value) for the driver mutation status term.

In the case of *TP53*, *IDH1*, and *PIK3CA*, each driver genes’ hits are also statistically enriched in known cancer-related genes, such as those from the Cancer Gene Census (CGC)^33^ (Fig. 2D, Table 1), highlighting Dyscovr’s ability to prioritize genes with cancer-relevant roles. This is further supported by gene set enrichment analysis: we find that our driver genes’ targets are also statistically enriched in cancer-related pathways from Gene Ontology (GO)^34^ and KEGG^32^ (Fig. S1C; Data S2). For *TP53*, for example, KEGG pathways include cell cycle (*q* = 7.84E-09), transcriptional misregulation in cancer (*q* = 1.82E-03), and p53 signaling pathway (*q* = 2.19E-03), as well as various metabolic pathways (Data S2).

**Table 1.** Top predicted dysregulated cancer genes from the Cancer Gene Census (CGC)^33^ in relation to mutations in driver genes. Ten most statistically significant (ranked by *q*-value) CGC dysregulated target genes from phase one of Dyscovr for each driver gene are shown, with those genes that are upregulated in relation to driver gene in orange and those that are downregulated in relation to driver gene in blue (pan-cancer).

| Driver Gene | 10 Most Significant Associated CGC<sup>5</sup> Genes |
| --- | --- |
| *TP53* | *MDM2*, *CDKN1A*, *DDB2*, *BUB1B*, *CDKN2A*, *FANCD2*, *STIL*,<br>*CHEK2*, *XPC*, *KNSTRN* |
| *PIK3CA* | *PIK3R1*, *ETV4*, *AR*, *TET2*, *BAZ1A*, *IDH2*, *SBDS*, *VTI1A*, *SOX2*,<br>*NOTCH2* |
| *KRAS* | *ETV4*, *ETV5*, *TCF7L2*, *CANT1*, *RHOA*, *SLC45A3*, *CREB3L1*,<br>*ARHGAP26*, *CEBPA*, *PPP2R1A* |
| *IDH1* | *TCF12*, *FBXW7*, *PRKCB*, *ID3*, *GAS7*, *MAML2*, *CHIC2*, *PRDM2*,<br>*HIF1A*, *TET2* |

Though an advantage of Dyscovr is its ability to estimate transcriptional changes in putative transcriptional targets individually, we find that many of the targets that Dyscovr prioritizes also interact with one another. When each driver gene’s top hits are overlaid on the STRING functional protein-protein interaction network^35^, subclusters of dysregulated processes emerge. In the case of *TP53*’s top hits, a cluster of interconnected, cell cycle-related genes linked directly to *TP53* or to one another are visible (Fig. 2E). Similar clusters can be observed for the other drivers, such as upregulated MAPK signaling in the case of *KRAS* and downregulated CaMK kinase cascade signaling in the case of *IDH1* (Fig. S2), suggesting that Dyscovr’s results can be used to visualize how driver gene mutations disrupt broad functional networks.

When top hits for each of these drivers are examined individually, known functionally relevant genes are apparent (Fig. 2F). These include *TP53* and *MDM2*, which form an autoregulatory feedback loop^36^, and *KRAS* mutation and upregulation of *DUSP4/6* and *ETV4/5*, which are members of the ERK/MAP cascade downstream of *KRAS*^37^. Another example is *PIK3CA* mutation and upregulation of *PIK3R3*, a regulatory subunit of the PI3-kinase (PI3K) of which *PIK3CA* is also a member. Dyscovr also identifies novel relationships with partners outside of the drivers’ immediate pathways, such as *PIK3CA* mutation and upregulation of sphingolipid metabolism genes *SGPL1* and *SPTLC2*. This class of genes has sparked recent interest due to its relevance to cancer diagnosis and prognosis, as well as its potential to provide new antitumor targets^38^. These links, which are available for browsing on the Dyscovr website, present intriguing, previously unstudied gene-gene relationships.

### Benchmarking Supports Dyscovr’s Ability to Uncover Dysregulated Targets

To further evaluate the robustness of Dyscovr’s predicted driver-target relationships, we perform several additional benchmarking and control analyses. When *TTN*–a gene mutated at greater than 5% frequency in 22 of these 32 cancers due to its long length–is included in the model, Dyscovr finds only three hits at *q* < 0.01 (Fig. 2A, Fig. S1A). Additionally, when randomizing gene expression data, no driver gene has significant hits (*q* < 0.2), suggesting that Dyscovr’s hits reflect genuine biological signal (Fig. S1B, Methods IV.A). We also evaluate a variant of Dyscovr in which only “hotspot” mutations (for *KRAS, PIK3CA*, and *IDH1*) or nonsense mutations and Clinical Interpretation of Variants in Cancer (CIVIC)^39^-supported missense mutations (for *TP53*) are considered. We find comparable performance when restricting to well-characterized driver alterations as when aggregating all nonsynonymous mutations across genes, as judged by enrichment in CGC genes (Fig. S3A), DoRothEA targets (Fig. S3B), and *TP53*-specific target sets (Fig. S3C).

We also compare the first component of Dyscovr, which links mutant cancer driver genes to dysregulated target genes, to other methods with comparable aims. We re-implemented muTarget^13^ (a Mann-Whitney U (MWU)-based method) on our pan-cancer cohort, finding that MWU produces skewed *p*-values (74.2% of driver-target pairs significant at adjusted *p*-value < 0.01, Fig. S4A). When tested on *TTN*, MWU finds 11,577 of 16,458 genes (70.3%) to be significantly dysregulated at adjusted *p* < 0.01, a dramatic number of false positives. Additionally, we observe that MWU results are significantly enriched for immune cell-specific marker genes, likely deriving from changes in expression in tumor-infiltrating immune cells rather than neoplastic cells (Fig. S4B). This is particularly confounded for *IDH1*, which is known to promote changes in the tumor immune microenvironment^40^ and appears to dominate the MWU signal, highlighting the importance of correcting for the fraction of infiltrating immune cells. Across all drivers, the overlap between Dyscovr and MWU’s top genes is limited, particularly for *PIK3CA* and *IDH1* (Fig. S4C). Additionally, among the top 500 genes, for example, Dyscovr consistently displays a greater proportion of CGC and DoRothEA target genes (Fig. S4C). This is also true for *TP53*-specific target genes, such as curated *TP53* targets^29^ and Reactome P53 pathway genes^41^ (Fig. S4D). We also benchmarked Dyscovr against xseq, a hierarchical Bayesian method that predicts the probability that mutations influence gene expression^5^. The xseq method uses a gene interaction network as a prior and therefore does not output predicted effects on every target gene. Nonetheless, when restricting the output of Dyscovr and MWU to the set of genes tested by xseq (i.e. neighbors in the provided interaction network) and testing for enrichment in curated *TP53* targets, as well as those from TRRUST^30^, KEGG^32^, Reactome^41^, and hTFtarget^31^, Dyscovr is more effective than xseq at prioritizing known targets (Fig. S4E).

### Dyscovr Uncovers Driver-Associated Dysregulated Genes within 19 Cancer Types

To discover associations between nonsynonymously mutated driver genes and the expression of other genes in a specific cancer context, we next applied Dyscovr separately to each of the 19 TCGA cancer types with at least 75 samples possessing all data types. Each TCGA cancer type has its own unique set of cancer driver genes^27^ mutated in at least 5% of samples (Fig. S5A); as such, the landscape of mutated driver-dysregulated gene pairs identified varies by cancer type (Fig. 3A). While the absolute number of significant hits at a fixed *q*-value threshold is in part dependent upon the number of available samples, all 19 cancer types possess at least 10 significant hits at a *q* < 0.2 threshold (Fig. 3A, Data S4).

**Figure 3.**
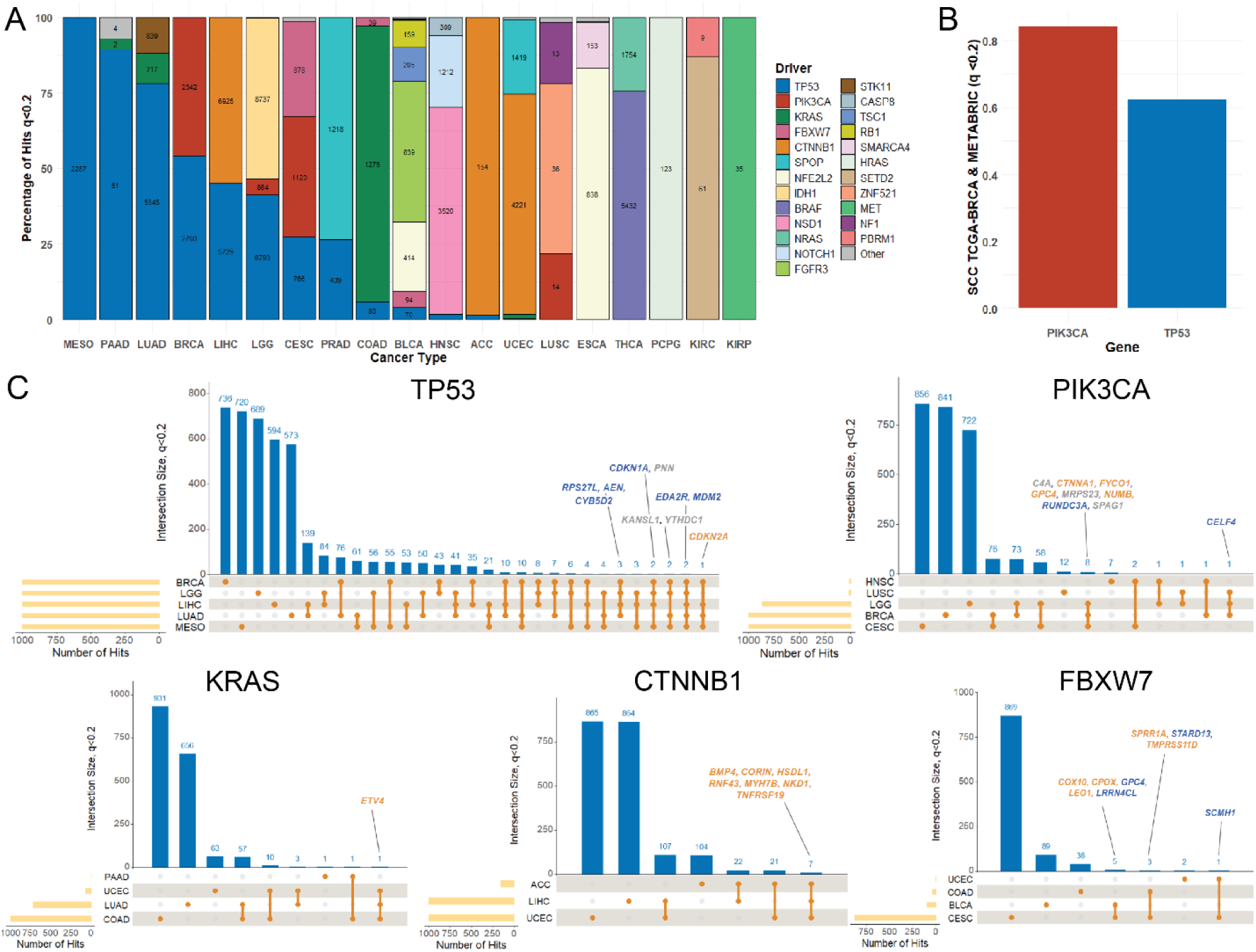
Dyscovr identifies associations between mutated cancer drivers and dysregulated gene expression across 19 TCGA cancer types. **A.** In each of 19 TCGA cancer types, the first stage of Dyscovr was applied to every candidate target gene, with each regression including covariates for all driver genes mutated at ≥5% frequency in that cancer type. Segments of the stacked bars represent the percentage of significant hits (*q* < 0.2) in each cancer type (*x*-axis) attributed to a given driver gene. Each segment is labeled with the absolute number of dysregulated targets associated with that driver in the corresponding cancer type (*q* < 0.2). See Table S1 for full cancer type names. **B.** Bar chart of the Spearman’s rank correlation (*y*-axis) between the nonsynonymous mutation status coefficient estimated by Dyscovr for the indicated drivers (*x*-axis, *PIK3CA* in red, *TP53* in blue) across target genes that are significant at *q* < 0.2 in two breast cancer cohorts: TCGA-BRCA and METABRIC. Pairwise Spearman correlation coefficients are 0.84 for *PIK3CA* and 0.62 for *TP53*. Hypergeometric enrichment *p*-values are 3.42E-06 for *TP53* and 1.50E-61 for *PIK3CA*. **C.** UpSet plots for the five driver genes with the largest absolute number of significant targets across at least three cancer types: *TP53, PIK3CA*, *KRAS, CTNNB1,* and *FBXW7*. Plots show the number of significantly dysregulated target genes (*q* < 0.2; capped at 1000 for visual clarity) in each of the up-to-five cancer types with the most hits for that driver (bottom left, yellow), as well as the intersections of targets shared across cancer types (blue). Significant targets shared among the most cancer types are labeled and colored by direction of regulation (upregulation in orange, downregulation in blue, variable direction in gray).

Interestingly, Dyscovr finds that transcriptional effects of driver mutations display similarity across cancer types, as the pairwise Spearman’s rank correlations of mutational coefficients fit by Dyscovr across all putative targets are largely positive (Fig. S5B). These results align with the hypothesis that mutations in cancer driver genes tend to affect a core set of downstream genes and processes across tissues. As expected, we still find a great deal of tissue specificity, however, as Dyscovr’s results more strongly replicate when the same tissue type is being compared. In the case of breast cancer, Dyscovr’s hits for TCGA-BRCA and external dataset METABRIC^42^ display high Spearman’s rank correlations (Fig. 3B, Fig. S5C) and overlap (HG *p* = 1.18E-07 and *p* = 2.45E-62, respectively) for the two drivers mutated in at least 5% of samples in both TCGA and METABRIC, *TP53* and *PIK3CA*. These hits represent genes that are commonly dysregulated across breast cancer subtypes, as subtype is included as a covariate in our regression (Methods IV).

We were particularly interested in targets that are recurrently dysregulated by the same mutated driver across multiple cancer types, as these mechanisms might be broadly targetable (Fig. 3C). Dyscovr identifies several such shared associations with known mechanisms-of-action; for example, activating *PIK3CA* mutations have been shown to result in upregulation of glypican (GPC) family members, such as *GPC4,* and consequent tumorigenesis in gliomas^43^. However, many recurrent associations identified by Dyscovr have not been previously described. For example, Dyscovr links *PIK3CA* mutations to *FYCO1* overexpression in breast cancer, cervical cancer, and low-grade glioma, an association that remains relatively unexplored despite evidence that *FYCO1* has important roles in migration and invasion of tumor cells^44^.

### Dyscovr Highlights Putative Genetic Interactions for TP53, PIK3CA, and KRAS

After identifying genes whose expression is associated with driver gene mutation status, Dyscovr evaluates these driver-target relationships to identify genetic interactions and prioritize conditional genetic vulnerabilities. Dyscovr uses a regression framework to analyze CRISPR dependency, somatic mutation, and gene expression data from the DepMap database^26^, with a single model per driver-target pair. This model tests whether driver nonsynonymous mutation status and gene expression level, both of which serve as proxies for driver activity, are associated with cell viability following CRISPR-Cas9 knockout of the target gene, while accounting for cancer type (Fig. 1D, Fig. 4A, Methods VIII). If no genetic interaction exists, the effect of target-gene knockout on cell viability should not depend on driver state. For each driver-target pair, evidence from the driver mutation and expression terms is combined using empirical Brown’s method to obtain a single *p*-value, followed by multiple-hypothesis correction (Methods VIII). Here, we apply this model to each of our TCGA pan-cancer driver genes with sufficient mutational diversity across CCLE cell lines (*TP53*, *PIK3CA*, and *KRAS*) and, for each driver, to the set of target genes found to be significantly dysregulated by that driver both pan-cancer and within at least one individual cancer type (*q* < 0.2). Using these criteria, Dyscovr systematically identifies hundreds of genetic interactions (Fig. 4B, Table S4, Data S5).

**Figure 4.**
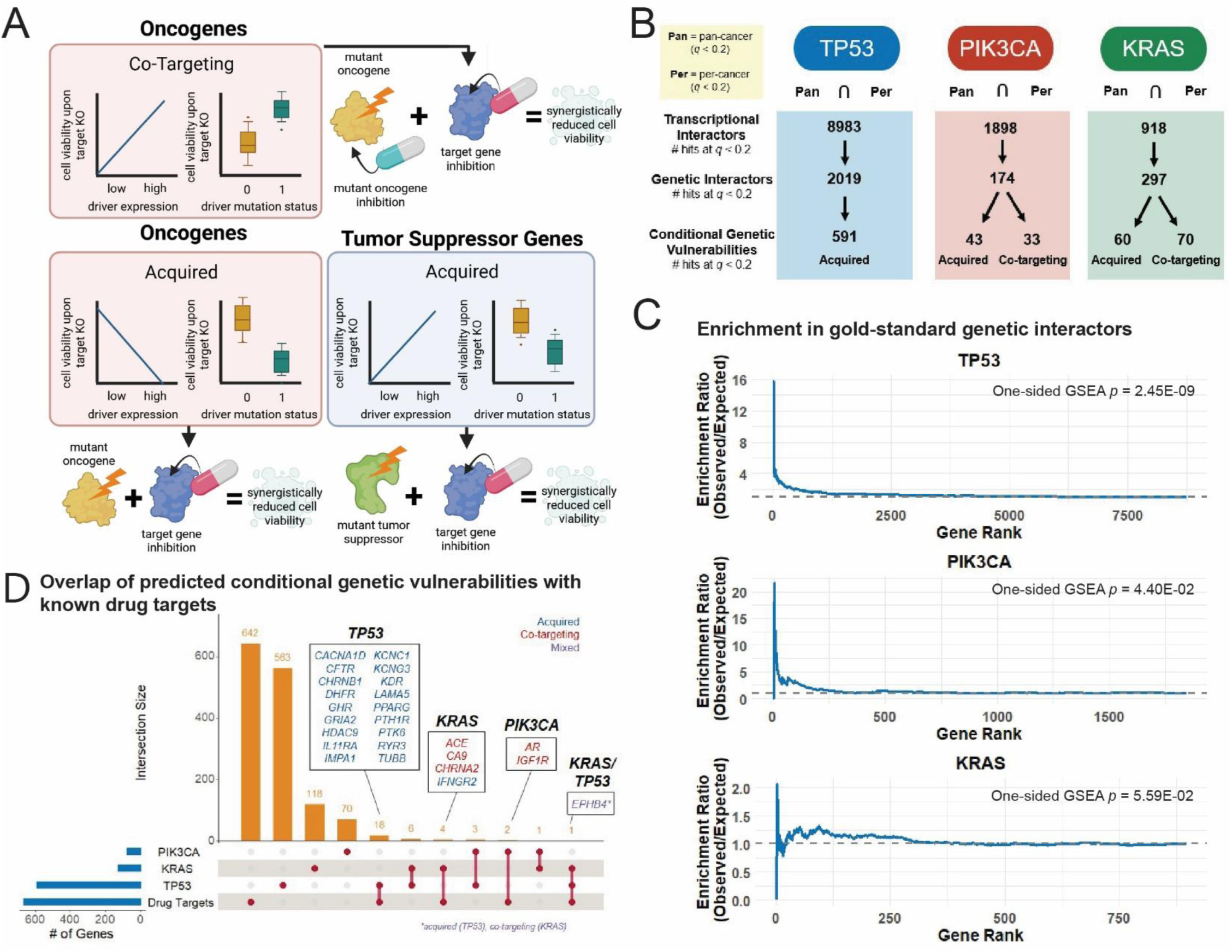
Dyscovr reveals genetic interactors with cancer driver genes *TP53*, *PIK3CA*, and *KRAS*. **A.** Schematic illustrating clinically relevant types of genetic interactions identified by the second stage of Dyscovr. For tumor suppressor genes (blue, i.e. *TP53*), Dyscovr identifies acquired vulnerabilities, in which mutations in the driver gene render cancer cells more sensitive to inhibition of the target gene. For oncogenes (pink, i.e. *KRAS* and *PIK3CA*), Dyscovr identifies both acquired and co-targeting vulnerabilities, where joint inhibition of the driver and the target gene more effectively reduces cell viability relative to inhibiting either gene alone. **B.** Schematic illustrating how Cancer Dependency Map (DepMap)^26^ cell line data and Dyscovr’s transcriptional predictions are integrated to infer putative genetic interactions. For pan-cancer driver genes with sufficient cell line representation (*TP53*, *PIK3CA*, and *KRAS),* we report the number of significant transcriptional hits from phase one of Dyscovr (*q* < 0.2) that are found both pan-cancer (“Pan”) and in at least one individual cancer type (“Per”) at the top. In the middle, we report the number of significant genes that are predicted to be genetic interactors with the associated driver, defined as genes whose knockout-associated cell viability is associated with driver activity. At the bottom, we report the number of significant genes classified as vulnerabilities either when the driver is mutated (“acquired”) or when jointly inhibited with the driver (“co-targeting”). **C.** Cumulative enrichment ratio of gold-standard genetic interactions (Methods VIII.B, Table S5) amongst Dyscovr’s ranked genetic interactors for *TP53* (gold-standard interactions N = 856), *PIK3CA* (N = 195), and *KRAS* (N = 4576). **D.** Intersection of clinically relevant conditional genetic vulnerabilities for each of *TP53, PIK3CA,* and *KRAS* with known US FDA approved drug targets^74^ (Methods VIII.C, Table S6). Genes with an associated drug are annotated and colored according to whether they were predicted to represent acquired or co-targeting vulnerabilities.

We next demonstrate that Dyscovr’s predicted genetic interactions are enriched for previously reported genetic interactors. In particular, we tested Dyscovr’s predictions for enrichment in a set of genetic interactions comprised of BioGRID^45^ and manually curated genetic interactors from the literature (Fig. 4C, Table S5, Methods VIII.B). For each driver gene, ranking its predicted genetic interactions by combined *p*-value, we observe enrichment of gold-standard genetic interactors among Dyscovr’s predicted genetic interactions for all three drivers (Fig. 4C), with statistically significant enrichment for *TP53* and *PIK3CA*, as well as strong enrichment for *KRAS*.

Among its predicted genetic interactions, Dyscovr further identifies those with therapeutic relevance, distinguishing cases in which the cancer-associated driver state creates or strengthens a dependency on the target (acquired vulnerabilities) from cases in which driver activity appears to buffer the effects of target loss, suggesting a potential benefit of jointly inhibiting of the driver and target (co-targeting vulnerabilities). Because tumor suppressors and oncogenes contribute to cancer through different changes in activity, we identify and interpret vulnerabilities differently for these two driver classes. Loss of tumor-suppressor activity can arise from disruptive nonsynonymous mutations or reduced expression, whereas increased oncogenic activity can result from activating mutations or elevated expression. Because loss-of-function in tumor suppressor genes is generally difficult to target directly, we focus on acquired vulnerabilities. These are indicated by negative associations between tumor suppressor mutation status and viability upon target knockout, consistent with increased sensitivity to target loss in the mutant state, as well as positive associations between tumor suppressor expression and viability upon target knockout, consistent with increased sensitivity when tumor suppressor expression is low (Fig. 4A). For oncogenes, whose activity can often be inhibited pharmacologically, we consider both acquired and co-targeting vulnerabilities. Acquired vulnerabilities are indicated by negative associations between oncogene activity and viability upon target knockout, whereas co-targeting vulnerabilities are indicated by positive associations between oncogene activation and viability upon target knockout, consistent with oncogene activity buffering the consequences of target loss (Fig. 4A).

Dyscovr identifies hundreds of predicted genetic interactions that correspond to these classes of conditional genetic vulnerabilities (Fig. 4B). We compare these vulnerabilities with interactions catalogued and labeled as SLs in SynLethDB^46^, a database focused cancer-associated SL relationships, as SL interactions constitute one of the best-studied forms of genetic vulnerability in cancer. For *TP53,* SynLethDB includes SLs in *TP53*-mutant contexts, corresponding to acquired vulnerabilities, as well as those in *TP53*-wild-type contexts, in which the presence of functional *TP53* increases dependency on the interacting gene, as illustrated by *MDM2*. Because SynLethDB but does not annotate the *TP53* context in which each interaction occurs, we test its entries for enrichment in a set consisting of Dyscovr’s predicted acquired vulnerabilities as well as its predicted interactions in which greater *TP53* activity is associated with increased sensitivity to target loss. For the oncogenes *PIK3CA* and *KRAS*, we focus on Dyscovr’s predicted co-targeting vulnerabilities, which most closely correspond to the simultaneous gene inhibition relationships catalogued in SynLethDB for these genes. Restricting Dyscovr’s predictions to these corresponding interaction classes, we observe enrichment for SLs supported by experimental or text mining-based evidence for all three drivers, with statistically significant enrichment for *TP53* (one-sided GSEA, NES = 1.71, *p* = 7.05E-05) and positive but non-significant enrichment for *PIK3CA* (NES = 1.26, *p* = 2.61E-01) and *KRAS* (NES = 1.15, *p* = 2.12E-01). We next consider the landscape of computationally predicted SLs included in SynLethDB, which highlights a major gap that Dyscovr helps address. Existing computational methods have made strikingly few predictions for the oncogenes *PIK3CA* and *KRAS*: only one gene, *EGFR*, is predicted by any method in SynLethDB to be SL with *KRAS*, a major limitation for an increasingly druggable oncogene^47^. Similarly, only 21 genes were computationally predicted to be SL with *PIK3CA* by any method. In contrast, SynLethDB contains 379 computationally predicted SLs for *TP53*, and we observe that predicted conditional vulnerabilities from Dyscovr are significantly enriched for these computationally predicted SLs (NES = 1.27, *p* = 3.20E-02). Together, these results suggest that Dyscovr not only can recapitulate known interactions but also substantially expands the landscape of predicted vulnerabilities for oncogenes that have been underserved by existing computational approaches.

Closer examination of the putative genetic interactors from Dyscovr reveals both known pairings from the literature and novel pairings, spanning both acquired and co-targeting vulnerabilities. The top co-targeting hit for *KRAS* is fibroblast growth factor receptor (*FGFR)* adaptor protein *FRS2*; *FGFR1*, which signals via *FRS2*, has been found to mediate adaptive drug response in *KRAS*-mutant lung cancers, leading to success of a combinatorial treatment approach^48^. Notably, Dyscovr identified this link between *KRAS* and downregulation of *FRS2* in lung cancer, but not in colon, pancreatic, or uterine: this is in close alignment with previous evidence that this combinatorial treatment strategy is effective only in *KRAS*-mutant lung cancers and not other *KRAS*-mutant cancer types^49^. Similarly, one of Dyscovr’s top acquired vulnerabilities for *KRAS* is *DUSP4*, which is a known SL gene of *KRAS*^50^. Other of *KRAS’s* top predicted genetic interactors have also been linked to *KRAS* mutations in a cancer context, such as predicted acquired vulnerability *IL17RE*^51^ and predicted co-targets *ARFGEF2*^52^ and *NPAS2*^53^.

In the case of *TP53*, the top predicted acquired vulnerability is *USP28*, an oncogene that regulates a variety of tumorigenic processes in cancers like squamous cell carcinoma, including cellular proliferation, DNA damage repair, and apoptosis. Overexpression of *USP28* has been associated with poorer outcomes, leading to the development of *USP28-*targeting therapeutics^54^. However, *USP28* has also been put forth as a candidate tumor suppressor, as its deubiquitinating functions play a role in stabilizing tumor suppressor *TP53 in vivo*^54^. Our analysis aligns with this dual functionality, suggesting that targeting *USP28* is most effective in contexts where *TP53* function has already been disrupted via nonsynonymous mutation and only *USP28’s* oncogenic roles remain relevant. Several others of *TP53*’s top predicted genetic interactors have evidence supporting an acquired vulnerability, including oncogene *CHEK2*^55^ (modulates resistance to epirubicin in tandem with *TP53* in breast cancer^56^); *LBR* (promotes cellular proliferation in the absence of *TP53*^57^); *ACO1/IRP1* (involved in a well-characterized iron-*TP53* feedback loop in cancer^58^); and *CUL9* (promotes *TP53*-dependent apoptosis^59^). Many of *TP53*’s other top hits have been reported in the cancer literature, though without a mechanistic relationship to *TP53* mutations, such as *NCDN*^60^, *PSME4*^61^, *FAM189*^62^, *CKAP2L*^63^, *GHR*^64^, and *MYO9B*^65^.

Many of mutant *PIK3CA*’s top predicted genetic interactors have been shown to act via or downstream of the PI3K-AKT signaling pathway, such as predicted co-targets *TM4SF1* (regulates breast cancer cell migration and apoptosis^66^); *SLC7A2* (mediates recruitment of myeloid-derived suppressor cells and tumor immunosuppression^67^); and *TCF7L2* (WNT pathway effector shown to mediate colorectal cancer cell migration and invasion^68^), as well as predicted acquired vulnerabilities *MOB1A* (Hippo pathway member that promotes PI3K-AKT signaling^69^) and *MADD* (a key cancer survival factor that is phosphorylated by *AKT*^70^). Other identified hits are involved in insulin signaling and regulation, which activates PI3K-AKT signaling; these include well-described *PIK3CA*-interactors *IRS2* and *IGF1R*^71^, which are also candidate oncogenes in a variety of cancer types^72,73^.

Each of mutant *TP53*, *PIK3CA*, and *KRAS* possesses genetic interactors predicted by Dyscovr that are targeted by US FDA approved drugs^74^ (Fig. 4D, Table S6). Certain drugs targeting acquired gene vulnerabilities are known to have improved effects in the context of driver mutation, such as pazopanib, which targets *VEGFR2/ KDR* and has greater efficacy in *TP53*-mutant sarcomas relative to those with wild-type *TP53*^75^. Additionally, some of Dyscovr’s predicted vulnerabilities possess experimental drugs that are effective in combination with drugs targeting the driver gene, such as the combination of alpelisib (targets mutant *PIK3CA*) and bicalutamide (targets *AR*)^76^ and LY294002 (targets PI3K) and OSI-906 (targets *IGF1R*)^77^. Altogether, the Dyscovr pipeline nominates a set of genes whose inhibition may either exploit dependencies associated with driver mutations or enhance the efficacy of therapies targeting oncogenic drivers, providing a compendium of candidate interactions with potential clinical actionability.

### Dyscovr Reveals Genes Co-targetable with Mutant Oncogenes

As a proof-of-concept demonstration of Dyscovr’s ability to identify candidate genetic interactors with potential clinical utility, we focused on *PIK3CA*, which possesses several inhibitors (including FDA-approved alpelisib and mutant-selective RLY-2606^78^). Among *PIK3CA*’s top ten predicted co-targeting partner genes from Dyscovr (Table S7), we selected an under-studied member of a cullin3-RING E3 ubiquitin ligase complex, kelch repeat and BTB domain-containing protein 2 (*KBTBD2)* for further study. Kelch repeat and BTB domain-containing proteins are adaptors which provide substrate specificity to the E3 ligase complex^79,80^. Physiologically, *KBTBD2* has been shown to regulate insulin signaling in adipocytes by controlling stability of PI3K regulatory subunit, p85α^81,82^. The function of *KBTBD2* in cancer, however, remains unexplored, though it was predicted by Dyscovr to be downregulated in relation to *PIK3CA* mutation pan-cancer (*q* = 6.5E-04) and in breast cancer (*q* = 4.8E-02), suggesting a cancer-relevant role. One possible explanation for this pattern is that *KBTBD2* participates in a negative feedback loop regulating PI3K signaling: under normal conditions PI3K activity promotes *KBTBD2* expression, while *KBTBD2* in turn limits pathway signaling through regulation of the PI3K regulatory subunit p85α. Constitutive activation of PI3K through oncogenic *PIK3CA* mutation may therefore suppress *KBTBD2* expression, disrupting this feedback control (Fig. 5A). Further implicating *KBTBD2* as a positive regulator of PI3K signaling in human cancer, co-dependency analysis showed that cells with a high dependence on *KBTBD2* were most likely to be sensitive to a variety of *IGF1R* inhibitors (Fig. 5B). Highlighting *KBTBD2* as a potential oncogene, we observed a significant reduction in survival rates across tumors in which *KBTBD2* was highly expressed, both pan-cancer (Fig. 5C) and in breast cancer (Fig. 5D), with cancer subtypes used as covariates in both analyses (Methods IX).

**Figure 5.**
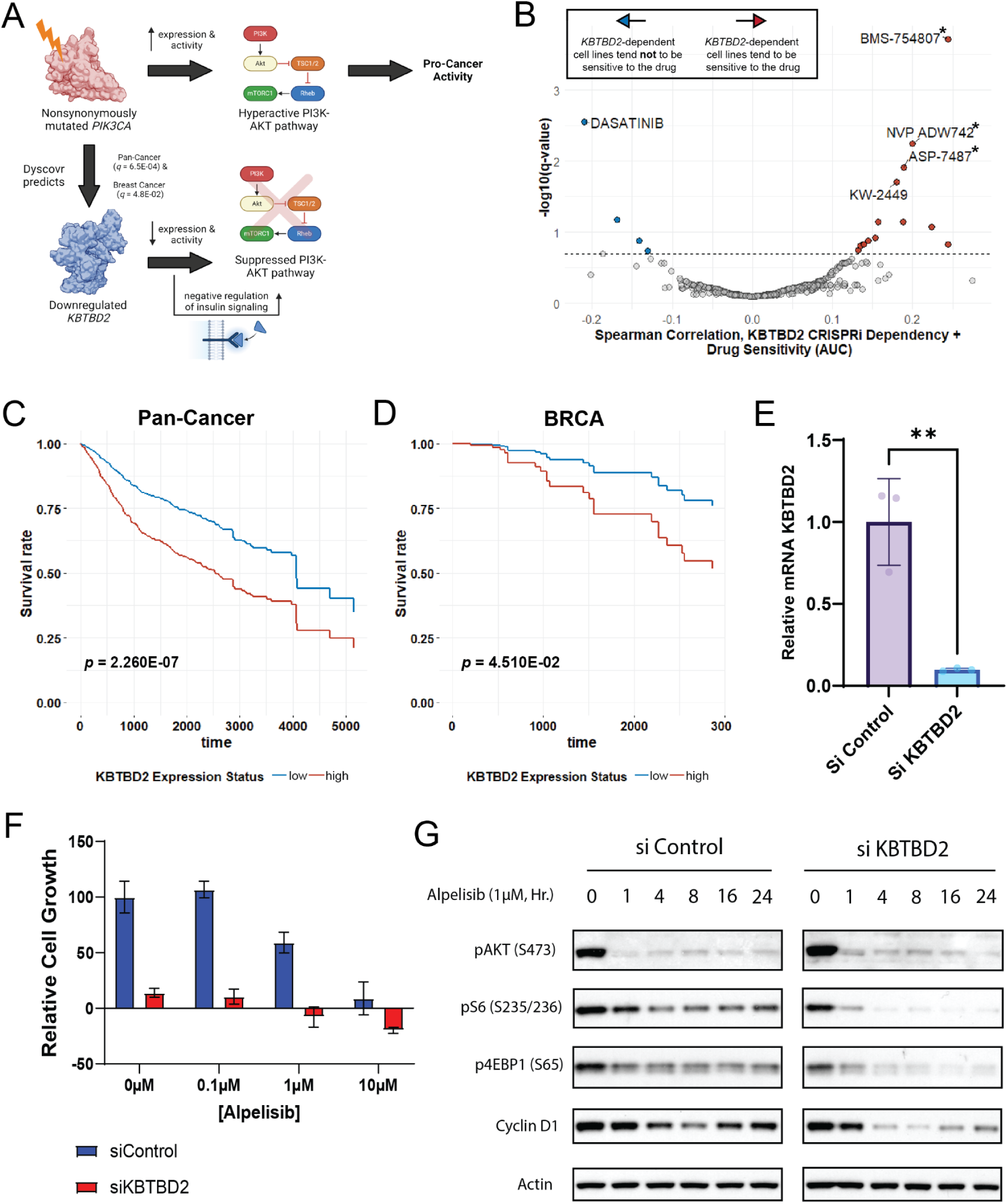
Mutant *PIK3CA* and *KBTBD2* expression exhibit a genetic interaction *in vitro*. **A.** Schematic illustrating a proposed mechanism-of-action underlying the predicted genetic interaction between mutated *PIK3CA* and dysregulated expression of *KBTBD2*. Dyscovr identifies an association between *PIK3CA* nonsynonymous mutation status and *KBTBD2* downregulation, both pan-cancer (*q* = 6.53E-04) and in breast cancer (*q* = 4.80E-02). This pair is also predicted to be a “co-targeting” vulnerability via cell viability analysis in the second phase of Dyscovr (*q* = 6.10E-02, Methods VIII.D). *KBTBD2* has been previously identified as being a regulator of insulin signaling in mouse models via its regulation of p85α protein abundance, with low levels of *KBTBD2* corresponding to suppressed PI3K-AKT signaling^81^. This proposed system suggests that inhibition of mutant *PIK3CA* may lead to recovery of *KBTBD2* expression; consequently, simultaneous inhibition of mutant *PIK3CA* and *KBTBD2* may produce synergistic negative effects on cell growth. **B.** The per-drug Spearman’s rank correlation between reported drug sensitivity scores from DepMap (AUC) and cell viability upon CRISPR knockout of *KBTBD2* (*x*-axis), plotted against the significance of that correlation (*y*-axis). Correlations were computed pan-cancer across cell lines and each of 4,659 drugs from DepMap. Drugs with statistically significant correlations to cell viability upon *KBTBD2* knockout (*q* < 0.2) are colored, with positive correlations in red and negative correlations in blue. Labels with an asterisk (*) denote drugs that target insulin like growth factor 1 receptor (*IGF1R*), suggesting that cell lines most sensitive to *KBTBD2* knockout also tend to be most sensitive to inhibitors of insulin signaling. **C-D.** Survival of pan-cancer (**C**) or breast cancer (**D**) TCGA patients significantly differs when stratified by “low” or “high” *KBTBD2* expression (*p* = 2.26E-07 and hazard ratio of 2.61 pan-cancer, *p* = 4.51E-02 and hazard ratio of 2.99 in BRCA). Survival rate (*y*-axis) over time in days (*x*-axis) was calculated using a Cox proportional hazards model that accounts for the nonsynonymous mutation status of *PIK3CA* and other clinical and molecular confounders (Methods IX). **E.** qPCR-based quantification of *KBTBD2* mRNA expression in MCF7 cells treated for 48 hours with an siRNA control or siRNA against *KBTBD2*. Data are presented as mean ± SD, and statistical significance is indicated (\*\**p* < 0.01). **F.** Relative cell growth in response to alpelisib treatment in siControl and siKBTBD2 treated cells. MCF7 cells were transfected with siControl or siKBTBD2 for 48 hours, followed by alpelisib treatment at indicated doses for an additional 2 days. Cell growth was quantified by SRB staining. Data are presented as mean ± SD. **G.** Western blot analysis of PI3K signal transduction following alpelisib treatment and *KBTBD2* knockdown. MCF7 cells were transfected with siControl or siKBTBD2 for 48 hours, followed by treatment with 1 µM alpelisib for time *t* (hours).

To interrogate the function of *KBTBD2 in vitro*, we used siRNA to ablate its expression in the ER+, PI3K E545K mutant breast cell line, MCF7. Knockdown of *KBTBD2* alone resulted in a 75-85% reduction in cell growth (Fig. 5E, Fig. S6A). Interestingly, knockdown of *KBTBD2* at baseline did not alter signaling through the PI3K pathway as analyzed by canonical substrates (Fig. S6B), though we did notice a slight increase in p85α expression–a regulatory subunit of PI3K involved in modulating insulin sensitivity–following ablation of *KBTBD2*. Strikingly, *KBTBD2* knockdown significantly enhanced the effects of PI3Kα inhibitor alpelisib^83^—with 1µM alpelisib blocking growth by approximately 25%, and the addition of *KBTBD2* ablation converting this effect into cytotoxic cell death (Fig. 5F). This effect was even more pronounced with 10µM alpelisib. To examine the effects on pathway output, we knocked down *KBTBD2* in MCF7 cells followed by treatment with alpelisib for time *t*. We analyzed phosphorylation of PI3K effector *AKT*, as well as the downstream output of mTOR complex I (*mTORC1*), which is regulated by PI3K signaling and is a frequently activated and therapeutically targeted oncogene across many cancer types^84^. As expected, suppression of *mTORC1* substrate phosphorylation correlates with response to PI3K inhibitors^82^. When *KBTBD2* was knocked down, we observed a deeper inhibition of *mTORC1* targets, *pS6* and *p4EBP1*. Cyclin D1, also under *mTORC1* control^85^, was also better suppressed when *KBTBD2* expression was ablated (Fig. 5G). We also tested the mutant-selective PI3K alpha inhibitor, RLY-2608^86^. In this case, *KBTBD2* knockdown was more effective at blocking growth than the drug at any concentration. Additionally, the combination of RLY-2608 with *KBTBD2* knockdown produced only a minor additive effect on growth inhibition. Interestingly, in terms of pathway inhibition, knockdown of *KBTBD2* enhanced the inhibitory effects of RLY-2608, with *pS6*, *p4EBP1*, and cyclin D1 all more deeply suppressed than in the control siRNA cells (Fig. S6C). Taken together, these results demonstrate that *KBTBD2* inhibition holds exciting potential to enhance the effects of *PIK3CA* inhibitors in *PIK3CA*-mutant breast cancers, warranting further investigation. This example illustrates how Dyscovr can link driver mutation-associated dysregulation to functionally relevant genetic vulnerabilities and identify interactions that may be clinically informative or even pharmacologically actionable.

## DISCUSSION

Cancer driver mutations reshape cellular programs far beyond the altered gene itself, propagating transcriptional changes across the genome that can reveal or contribute to conditional vulnerabilities in tumor cells. Distinguishing therapeutically actionable vulnerabilities from expression changes that reflect broader features of tumor development (such as co-occurring driver alterations, widespread CNAs, or immune cell infiltration) remains challenging. Uncovering genes that can be co-targeted with oncogenic drivers, many of which are activated rather than lost, has proven especially difficult for existing genetic interaction discovery approaches.

Here, we introduced Dyscovr, a two-stage framework designed to address this gap. In the first stage, Dyscovr uses multiomic data from primary tumors to identify genes whose expression is systematically altered in association with nonsynonymous mutations in driver genes. By accounting for the diverse molecular and clinical factors that shape gene expression in tumors, Dyscovr is designed to distinguish expression differences associated with driver mutation status from those attributable to correlated features of the cancer context. Our results show that sources of variability such as cancer subtype, CNAs, DNA methylation, and immune cell infiltration are strong determinants of a target gene’s expression (Fig. S7A, Table S8), suggesting that attempts to relate mutation status to target expression without accounting for these factors may lead to misleading conclusions in the cancer context. When applied to more than 6,000 TCGA tumors across 19 cancer types, Dyscovr identifies hundreds of driver-associated transcriptional changes that are reproducible across patient cohorts and are enriched for known and cancer-related target genes; the linear modeling framework further facilitates interpretability of individual driver-target associations.

In the second stage, Dyscovr prioritizes these dysregulated genes using functional genetic screening data from cancer cell lines, distinguishing genes that become vulnerabilities in driver-mutant contexts from genes whose inhibition may enhance the effect of inhibiting the driver itself. Together, these two stages nominate genes that are both transcriptionally perturbed in association with a mutant driver and functionally relevant for the viability of cells harboring that alteration. Dyscovr highlights genes whose inhibition may either exploit a dependency associated with the driver mutation or provide a co-targeting strategy to enhance therapies directed at the driver, thereby nominating candidates for both single-agent and combination treatment strategies. Notably, Dyscovr can uncover genetic interactions for oncogenic drivers, for which classical synthetic lethal paradigms are less directly applicable. Many of the genes identified by Dyscovr as genetically interacting with driver genes possess targeted therapeutics in various stages of clinical development, opening possibilities for drug repurposing (Fig. 4D).

We experimentally validated a predicted genetic interaction between *PIK3CA* and *KBTBD2,* a gene our analysis implicates as a putative regulator of insulin and PI3K-AKT signaling. In *PIK3CA*-mutant breast cancer cell lines, joint inhibition of *PIK3CA* and *KBTBD2* more effectively suppressed cell growth than *PIK3CA* inhibition or *KBTBD2* knockout alone (Fig. 5). This finding illustrates how Dyscovr can nominate mechanistically interpretable interactions and reveal potential co-targeting strategies for oncogenic drivers. All genetic interactions predicted by Dyscovr are accessible via its website (dyscovr.princeton.edu), providing a resource for exploring putative interactions across genes and cancer types.

Several directions could further expand the Dyscovr framework. Our current analysis focuses on driver genes mutated at sufficient frequency across available tumor cohorts; as multiomic datasets continue to grow, the approach can be extended to rarer driver alterations and more genetically diverse tumor types. Similarly, while we currently account for tumor subtype as a confounder (Table S10), larger patient cohorts would provide statistical power to uncover subtype-specific interactions. Incorporating additional molecular features, such as chromatin accessibility or large-scale proteomic data, could also refine predictions by directly linking mutated driver genes to protein-level dysregulation, a more immediate therapeutic substrate compared to RNA. Finally, integrating patient survival data or clinical trial results could help narrow Dyscovr’s predictions to those with the greatest translational promise.

In sum, Dyscovr provides a systematic framework for linking mutations within cancer driver genes to transcriptional dysregulation and conditional genetic vulnerabilities. By connecting tumor multiomic data with functional dependency screens, Dyscovr nominates driver-associated genetic interactions for further experimental and therapeutic investigation. Because driver mutation profiles are now routinely obtained in clinical settings, Dyscovr’s predictions have the potential to inform therapeutic prioritization by connecting the growing scale of tumor genomic profiling to experimentally testable hypotheses that could support the development of more effective, driver-specific treatment strategies.

## ACKNOWLEDGEMENTS

Thanks to members of the Singh and Rabinowitz labs for their insights and comments. Thanks especially to Joshua Wetzel for helpful discussions regarding TCGA mutation and CNA data and for his review of the manuscript. The results published here are in part based upon data generated by the TCGA Research Network: http://cancergenome.nih.gov/, the METABRIC Consortium: https://www.nature.com/articles/nature10983, and the DepMap Consortium: https://depmap.org/portal/. This work was funded in part by NIH grant R01-CA208148 (to M.S.), Ludwig-Princeton AGMT DTD 1/1/2021 (to J.D.R. and M.S.), and NSF GRFP grant DGE-2039656 (to S.G.). The funders had no role in study design, data collection and analysis, decision to publish, or preparation of the manuscript. Original figures created using BioRender.com.

## AUTHOR CONTRIBUTIONS

Conceptualization, S.G. and M.S. (Dyscovr), J.B., M.M., and J.D.R. (Experimental Validation); Methodology, S.G. and M.S.; Software, S.G. (Dyscovr), N.A. and L.P. (Dyscovr Website); Validation, S.G. (Computational), J.B., M.F., and R.R. (Experimental); Resources, M.S. and J.D.R.; Writing – Original Draft, S.G.; Writing – Review & Editing, S.G., M.S., J.B., and J.D.R.; Visualization, S.G. and J.B.; Supervision, M.S. and J.D.R.; Project Administration, M.S. and J.D.R; Funding Acquisition, M.S. and J.D.R.

## METHODS

### I. A Framework to Estimate Regression Coefficients for the Nonsynonymous Mutation Status of Driver Genes

We introduce a linear regression framework to estimate relationships between the mutation status of driver genes and the expression of each putative target gene, *t*, across a set of cancer samples. In particular, for each putative target gene *t*, we consider the following model:

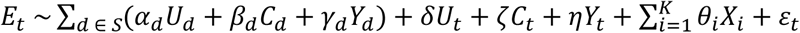

where *S* is the set of frequently mutated driver genes considered (Table S1, Methods IV.E), *E*_*t*_ is a continuous value representing the expression of putative target gene *t* (Methods IV.A), *U*_*d*_ is a binary variable indicating whether driver gene *d* possesses a nonsynonymous mutation (Methods IV.B), *C*_*d*_ is a continuous value representing the normalized copy number status of *d* (Methods IV.C), and *Y*_*d*_ is a continuous value representing the methylation status of *d* (Methods IV.D). Similarly, *U*_*t*_ is a binary variable indicating whether the putative target gene *t* possesses a nonsynonymous mutation, *C*_*t*_ is a continuous value representing the normalized copy number status of *t,* and *Y*_*t*_ is a continuous value representing the methylation status of *t.* In addition to these core features, we also have a number of additional covariates that correspond to various other clinical and molecular features that may have nontrivial effects on *E*_*t*_, with the number *K* of such covariates dependent upon the characteristics of the set of samples being examined (Methods IV.F). These covariates include the cancer type and subtype, age, sex, genotypic background, prior malignancies, prior treatment, nonsynonymous tumor mutational burden (TMB), tumor purity, and fraction of infiltrating immune cells. The coefficients fit from the data are the α_*d*_, β_*d*_, γ_*d*_ and {θ_1_, θ_2_, …, θ_*K*_}, as well as δ, ζ, and η.

### II. Applying Regression Framework to Samples from the TCGA, Pan-Cancer and Within 19 Individual Cancer Types

We apply the multiple linear regression model to all putative target genes *t* in the human genome (Methods IV.A), with one model per *t*. We do this both across all TCGA samples, or “pan-cancer”, as well as within each of 19 individual TCGA cancer types possessing ≥75 samples (Table S1). For both the pan-cancer analysis and the per-cancer analyses, each model includes *U*_*d*_, *C*_*d*_, and *Y*_*d*_ features for all driver genes *d* as annotated by Vogelstein et al.^27^ that have nonsynonymous mutations in at least 5% of available samples and at least 5 total samples; see Table S1 for the set of drivers tested in each case. In the pan-cancer case, drivers must be mutated in at least 5 samples, as well as at least 5% of all samples both pan-cancer and within at least 2 individual cancer types (to ensure signal is not driven by a single cancer type). This 5% threshold was chosen in an effort to balance statistical power with including as many potentially interesting driver genes as possible. All multiple regression models were run using the R package speedglm’s *speedlm* function^87^, which fit coefficients by minimizing the residual sum of squares and provided associated *p*-values for all terms, including the *U*_*d*_ mutational terms of interest. For all tests, *p*-values are two-sided and correspond to the F-statistic, which is calculated according to the null hypothesis that a given coefficient is zero. For each driver gene, we performed multiple hypothesis correction across the set of coefficients corresponding to its mutational term to convert *p-*values across the targets to *q*-values, using the *qvalue* function from the qvalue package in R with default parameters^88^. We deemed pairings between a nonsynonymous mutation in driver gene *d* and the expression of putative target gene *t* significant if the corresponding *q*-value was less than a threshold value of 0.2; given the increased statistical power in the pan-cancer setting, in the main body of the paper, we report pairings that are significant using a threshold of 0.01 for pan-cancer analyses and 0.2 for per-cancer analyses.

### III. Addressing Multicollinearity

In certain cases, we expect the nonsynonymous mutation status of driver gene *d*, represented as *U*_*d*_, to result in correlations with other variables in our framework (e.g., mutations within *IDH1* promote hypermethylation^89^, and thus *IDH1* mutation status in some cancers may be correlated with the methylation status of genes). In such cases, we will not be able to disentangle the contributions of the mutation within the driver gene from other variables it is correlated with. As such, prior to running the regression, our framework checks for multicollinearity.

First, we check that cancer subtype covariates (Methods IV.F.h) are not correlated with the mutation status of any driver genes present in the given regression model. If they are, this would indicate that these subtypes were at least partially defined by driver mutation status. To address this, we generate a Spearman correlation coefficient matrix and associated *p*-value matrix using the R package Hmisc, and check, for each cancer subtype variable, if it is correlated with any *U*_*d*_ variable with an absolute Spearman correlation >0.7 and *p*-value <1E-05. If this correlation meets both of these exclusion criteria, the subtype variable is removed from the regression model. We use Spearman correlation for subtype variables, rather than other commonly used measures of multicollinearity such as the variance inflation factor (VIF)^90^, because subtypes are encoded as bucketed, binary variables (Methods IV.F.h), and thus the corresponding variables are correlated with each other and will have high variance inflation factors (VIFs), even if they are not correlated with the mutation status of the driver gene. Across targets, we find that the vast majority of variables removed using this procedure pan-cancer (∼99.9%) correspond to subtypes that are defined by IDH1-mutation status in LGG, particularly the IDH1mut-non-codel subtype.

Following this, we check other variables in the model for multicollinearity using the VIF. We use the R package caret’s *vif* function to calculate a VIF measure for all non-bucketed variables in the regression, since as described above, bucketed variables (Methods IV.F.f-i) by design are collinear with one another and have high VIF scores. For the remainder of these variables, we eliminate any non-driver mutation (*U*_*d*_) variables whose VIF score exceeds a threshold of 5, a generally conservative but widely accepted threshold that suggests moderate collinearity^39^. To ensure that our variables of interest, *U*_*d*_, are not collinear with other variables in the model, we repeat this process iteratively until *U*_*d*_ for all *d* in *S* are below the threshold VIF of 5. In pan-cancer analyses, 13,917 genes (84.6%) had at least one variable removed, though 13,384 of these genes (96.2% of the 13,917 cases, 81.3% of target genes overall) had only the IDH1mut-non-codel subtype variable removed. Aside from this, we find that the methylation status of the target gene is most commonly eliminated pan-cancer (for ∼2.10% of tested genes), followed by sex (∼1.51%). For all significant pairings, variables that were eliminated using either of the above techniques are reported in Data S3 and Fig. S7B.

### IV. Data Acquisition and Processing

In the National Cancer Institute’s GDC data portal, there are 6378 primary tumor samples that possess all data types of interest, including annotated somatic mutation data, transcriptomic profiling, copy number variation, methylation, and clinical data. We downloaded these files from the GDC data repository, with parameters provided in Table S9.

A. *<u>Expression Data</u>.* Raw read count RNA-seq files from the TCGA were converted to counts per million (CPM) using edgeR’s *cpm* function^91^. Genes minimally expressed across all samples were eliminated using edgeR’s *filterByExpr* function with default parameters. These include requirements that all genes are required to have a minimum overall total count of at least 15 (*min.total.count*), and a minimum CPM of 10 in at least 10 samples (*min.count*). The remaining 19,052 genes’ counts were quantile normalized so that each of the 6378 samples has the same distribution of gene expression values using scikit learn’s *quantile.transform* function^92^, with an output distribution of ‘normal.’ Each gene *t*’s expression level in Dyscovr corresponds to a quantile-normalized gene expression value *E*_*t*_. For randomization analyses, each genes’ quantile-normalized expression values were shuffled across patients using the R function *dqsample*.
B. *<u>Mutation Data</u>.* We imported the simple nucleotide variation (SNV) file, with mutations called using *muse*^93^, into R using maftools’ *read.maf* function^94^. We then subsetted this file to include only nonsynonymous mutations (as annotated by *muse*), including missense, nonsense, nonstop, and splice site mutations. We then used maftools’ *mutCountMatrix* function to compute the number of nonsynonymous mutations per gene and per sample. We excluded samples with excessively high mutation rates across all genes, referred to here as ‘hypermutators’, which we defined according to the analyses performed in Campbell et al.^95^. In this work, they used a linear regression approach across 81,337 cancer patients to determine a reasonable threshold for hypermutation, which they recommend being ∼10 mutations per Mb. Given that the human exome is ∼36.8Mb, we selected 368 mutations as our threshold for hypermutation, such that any sample with greater than 368 total nonsynonymous mutations was discarded from the analysis. This removed a total of 360 samples, leaving 6018 pan-cancer TCGA samples. The mutation status of each gene *t* in each sample, *U*_*t*_, is 1 if the gene has a nonsynonymous mutation in that sample and 0 otherwise.
  a. <u>Restricting to Hotspot or Nonsense Mutations</u>. In some analyses, we restricted only to mutations that we expect to have high impact in terms of effect on protein function. For oncogenes *PIK3CA*, *KRAS*, and *IDH1*, we used literature-curated “hotspot” mutation residues. For *PIK3CA*, these include E542, E545, H1047 and H1047^96^. For *KRAS,* these include G12, G13, and Q61^97^. For *IDH1*, this includes R132^98^. For tumor suppressor gene *TP53*, we used all missense mutations with at least one line of evidence in Clinical Interpretation of Variants in Cancer^99^ and all nonsense variants (collectively 36.1% of *TP53* mutations). For analyses using these restricted mutation sets, we excluded patients with mutations in these drivers that lie only in non-hotspot positions.
C. *<u>Copy Number Alteration (CNA) Data</u>.* We obtained absolute CNA values for each gene in each sample, as computed by the *ASCAT*^100^ pipeline and made available by TCGA. In our model, for each gene in a given sample, we compute its normalized copy number value as the *log2* of the absolute CNA value divided by the mean CNA value across genes in the sample. Pseudocounts of 1 were used to adjust both the gene-level and average copy number values. The normalized copy number gives us a value that accounts for large scale ploidy differences between tumor samples. In practice, to be robust to extreme outlier CNA events, we exclude the top 10% and bottom 10% of gene-level CNA values for each sample when calculating its mean CNA value.
D. *<u>Methylation Data</u>.* We imported individual samples’ level 3 Liftover methylation Beta (β) files into R and removed ‘NA’ or empty values. These files have already been processed to include gene-level annotations. For any gene with more than one reported β value, we averaged these β values to produce a single β value per gene. Subsequently, we compiled the average β value for each gene in each sample and then converted β values to *M*-values using the logit, or *log*2(β / 1 − β)). We represent methylation levels in our model using these *M*-values. We use the *M*-value rather than the β value to assess methylation due to its purported statistical rigor^101^ and improved model performance in early testing.
E. *<u>Gene Set Data</u>.* Our list of driver genes consists of genes from Vogelstein *et al.* tables S2A, S2B, S3A, S3B, S3C and S4^27^.
F. *<u>Clinical Data</u>.* Using data from the TCGA’s clinical supplement, we created a composite data table with patient-level features. Using data from TCGA’s biosample data supplement, a TCGA tumor purity file obtained from Aran *et al.*^102^, genotypic principal components from Carrot-Zhang *et al.*^103^, and immune cell infiltration estimates from R’s immunedeconv package^104^ using the tool CIBERSORT Abs^105^, we created a data table with sample-level features. We then combined the patient– and sample-specific files such that all patient-specific data was applied to all samples from that patient, for use in the linear regression framework. Further information on each clinical feature is described in subsections below.
  a. *<u>Age.</u>* Normalized to take on a value approximately between 0 and 1 by dividing each patient’s age (given in years) by 100.
  b. *<u>Sex.</u>* Takes on a binary value of 1 for male and 0 for female patients.
  c. *<u>Prior malignancies.</u>* Takes on binary value, with 1 signifying that the patient had a prior malignancy and 0 signifying that there was no prior malignancy.
  d. *<u>Prior treatment.</u>* Encompasses two binary variables, as per the data available in the TCGA’s clinical supplement: prior radiation treatment, where 0 represents no prior radiation treatment and 1 represents prior radiation treatment, and prior pharmaceutical treatment, where 0 represents no prior pharmaceutical treatment and 1 represents prior pharmaceutical treatment.
  e. *<u>Genotypic Principal Components</u>*. Consists of three continuous covariates, each of which corresponds to the value of one of the first three genotypic principal components (PCs). These PCs are provided in the supplemental data table of Carrot-Zhang et al.^103^ (https://gdc.cancer.gov/about-data/publications/CCG-AIM-2020, “WashU_PCA_ethnicity_assigned.tsv”) and were calculated using the Washington University approach. Briefly, this approach involves the conversion of Birdseed genotype files to individual VCF files, which are then merged (with only variants of MAF >15% retained) prior to PCA using PLINK 1.9.^106^ See Carrot-Zhang et al.^103^ for more detailed methods.
  f. *<u>Nonsynonymous Mutational Burden.</u>* Consists of three binary covariates, representing low, moderate, and high nonsynonymous tumor mutational burden (TMB). For any given sample, one of these covariates will take on a value of 1 and the other two a value of 0, depending on the total number of nonsynonymous mutations their tumor possesses. Based on the distribution of TMB across TCGA samples, we defined low TMB as having <30 nonsynonymous mutations, moderate TMB as between 30 and 60 mutations, and high TMB as >60 mutations.
  g. *<u>Tumor Purity.</u>* Consists of three binary covariates, representing low, moderate, and high tumor purity. For any given sample, one of these covariates will take on a value of 1 and the other two a value of 0, depending on the magnitude of the tumor purity estimate for that sample. We define purity using the combined purity estimate (CPE) from Aran *et al.*^102^ when available, and for the samples without a provided CPE, we use the median of the available purity measures. Based on the distribution of CPE values across TCGA samples, we defined low tumor purity as having a CPE ≤ 0.5, moderate tumor purity as having a 0.5 < CPE ≤ 0.75, and high tumor purity as having a CPE > 0.75.
  h. *<u>Tumor Subtype</u>.* Consists of a binary covariate for each subtype in the given cancer type. For any given sample, one of these covariates will take on a value of 1 and all others a value of 0, depending on which subtype it is classified as. Because subtypes are defined differently for each cancer type, the column we used to define subtype in the TCGA clinical supplement is provided in Table S10. Each cancer type will have a different number of tumor subtype variables, as this is dependent on the number of unique subtypes present in the given column. Molecular subtypes were used whenever possible; when unavailable, histological or expression-based clustering subtypes were used. In the pan-cancer analyses, a binary covariate was created for each cancer type:subtype combination (e.g. Breast Cancer:Luminal A), across all cancer types. In this case, for any given sample, one of these combination covariates will take on a value of 1 and all others 0, depending on its combined cancer type and subtype classification.
  i. *<u>Immune Cell Infiltration</u>*. In the case of immune cell infiltration, we use the absolute immune cell fractions provided by the tool CIBERSORT Abs^105^, run using R’s immunedeconv package^104^. To get a single value representing the level of immune cell infiltration in the given sample, we add individual immune cell type fractions, e.g. predicted fractions of B cells, T cells, etc., into a total fraction of immune cells per sample. From there, we group this fraction into one of three buckets: low immune cell infiltration (total immune cell fraction ≤0.3), medium immune cell infiltration (total immune cell fraction >0.3 and ≤0.7), and high immune cell infiltration (total immune cell fraction >0.7), again defined by the distribution of total immune cell fractions across the samples.
G. In all of the cases that involve two or more binary covariates, and in which only one of these covariates can take on a value of 1 (Methods IVFf-i), one fewer covariate is needed in the linear regression equation to represent all distinct possible classifications^107^. To limit multicollinearity, the final covariate for a given feature is removed (i.e. a dummy variable encoding, rather than a one-hot encoding). For example, in the case of nonsynonymous TMB, low TMB would be represented by a binary covariate taking on values of 1 and 0, moderate TMB would be represented by a binary covariate taking on values of 0 and 1, and high TMB would be represented by two binary covariates taking on values of 0 and 0.

### V. Benchmarking of Dyscovr Against muTarget and xseq

We sought to benchmark Dyscovr’s performance in recovering transcriptional targets of driver genes as compared to muTarget^13^ and xseq^5^. In the case of muTarget and xseq, we performed benchmarking in a pan-cancer context and, for muTarget, used our preprocessed TCGA patient data as input (Methods IV). Each implementation is described in more detail below:

A. <u>muTarget (Mann-Whitney U).</u> Though muTarget did not provide unfiltered, downloadable results or associated source code, we re-implemented the underlying method using the *wilcox.test* function in R (with exact set to False), which performs the Wilcoxon rank-sum test (equivalent to the MWU). For each of *TP53*, *PIK3CA*, *KRAS*, and *IDH1*, we compared the expression of a given target gene between patient populations possessing a nonsynonymous mutation in that driver to those without, retrieving an associated *U*-statistic and *p*-value. We performed this test across all genes considered in the Dyscovr model, for a total of 16,458 tests. Given the high level of resulting *p*-value skew (Fig. S2A), *q*-value adjustment (which relies on an expected underlying *p*-value distribution) was not possible. Therefore, we adjusted *p*-values using a Benjamini-Hochberg correction. We ranked genes by *p*-value to examine overlap with Dyscovr’s hits, compare driver-target ranks, and perform gene set enrichment analysis (Methods VII).
B. <u>xseq.</u> We cloned the xseq v0.2.2 GitHub repository (https://github.com/shahcompbio/xseq) and downloaded the recommended influence graph (http://compbio-bccrc.sites.olt.ubc.ca/files/2015/05/influence_graph.txt), limiting it to connection strength greater than or equal to 0.4 as suggested. From there, we followed the provided vignette (https://rdrr.io/cran/xseq/f/vignettes/xseq-package.Rmd). Briefly, this involved converting our mutation, CNA, and expression files from the TCGA (Methods IV.A-C) into xseq format using the provided helper functions; imputing NA values in each matrix using K-nearest neighbor averaging, as suggested; obtaining the conditional distributions across genes; limiting to expressed genes using the recommended weight threshold of 0.9; setting model parameter priors; initiating the xseq model and learning the associated parameters; and converting the output into readable format. For the *trans* version of the xseq model, which is most comparable to Dyscovr, we followed the recommended protocol to remove *cis*-effects of CNAs on gene expression using the *NormExpr* function, and then ran *SetXseqPrior* and *InitXseqModel* with *cis* set to False. As output, we obtained combinations of drivers and target genes with an associated *P(D)* value, which corresponds to the conditional probability that a mutation in the given driver is associated with a change in expression in the given target gene. We used this descending probability for ranking our driver-target pairs. Given that only genes affiliated with a given driver in the provided influence graph possessed an associated *P(D)* value using this method (932 for *TP53*, 422 for *PIK3CA*, 213 for *KRAS*, and 54 for *IDH1*), we subsetted Dyscovr and muTarget’s output to the same genes for all subsequent enrichment tests.

### VI. Computational Validation via Comparison of TCGA BRCA results to METABRIC

To test whether our models are capable of generating consistent and meaningful correlations between driver gene mutations and target gene expression across independent cohorts, we also applied the Dyscovr framework to data from the Molecular Taxonomy of Breast Cancer International Consortium (METABRIC)^42^, a collection of 854 breast cancer patients of European ancestry with paired mutation, CNA, mRNA expression, methylation, and clinical data. Due to differences in available data types, we made the following modifications to our model and data processing.

A. <u>METABRIC: Estimating Regression Coefficient for Mutation Status of Driver Genes.</u> To ensure comparability to TCGA-BRCA, linear regression models were constructed in the same fashion as described in Methods I. Due to limitations in available germline data for the METABRIC cohort, genotypic principal components are not included in these models.
B. <u>METABRIC: Data Acquisition and Processing.</u> METABRIC primary tumor data files were downloaded from cBioPortal^108^. To keep processing pipelines as similar as possible to TCGA-BRCA, the same protocol was used for preprocessing mutation data, including generation of a mutation count, restriction to only nonsynonymous mutations, and removal of hypermutators (Methods IV.B). Mutations in METABRIC were called using MuTect and filtered according to the procedure given in Curtis et al.^42^. As with TCGA-BRCA, *ASCAT*^100^ was used for CNA calling. The gene expression data available for the METABRIC cohort is Illumina HT-12 v3 microarray data, which requires distinct preprocessing procedures to TCGA-BRCA’s RNA-sequencing data. We filtered genes with greater than 50% missing values and with mean expression <5 or standard deviation <0.3 across samples, as in Liao et al.^109^, but otherwise used the provided quantile-normalized *log2*-intensity values as input to our models. The METABRIC methylation data is also distinct from TCGA-BRCA in that only promoter methylation bisulfite sequencing (RRBS) is available. Files were already processed using the gpatterns package such that each file contains a [0,1] gene-level value of CpG methylation. Models were run using both these [0,1] CpG values, as well as the logit of these values, though models performed comparably and the [0,1] CpG values were ultimately used in figure creation. Due to differences in labeling and availability of clinical data types in METABRIC as compared to TCGA-BRCA, different column names were often used; any features with notable differences are described below.
  a. *Prior malignancies (Methods IV.F.c)*. The “RFS_STATUS” column was used.
  b. *Prior treatment (Methods IV.F.d).* The “RADIO_THERAPY” column was used to create a binary representation of prior radiation treatment, while the “CHEMOTHERAPY” and “HORMONE_THERAPY” columns were combined to create a binary representation of prior pharmaceutical treatment. In the latter case, the sample received a 1 if they had received either chemotherapy or hormone therapy, and 0 otherwise.
  c. *Nonsynonymous tumor mutational burden (TMB) (Methods IV.F.f).* For the nonsynonymous TMB, the *log2* of the “TMB_NONSYNONYMOUS” column with a pseudocount of 1 was used, with 3 binary covariates for representing low (TMB ≤ 2.5), moderate (2.5 < TMB ≤ 3.5) and high (TMB > 3.5) tumor mutational burden.
  d. *Tumor purity (Methods IV.F.g)*. METABRIC’s clinical supplement provides a “CELLULARITY” column that is an estimate of tumor purity from the tool MCP-counter^110^. The values this column can take include “Low”, “Moderate”, and “High”. As with TCGA-BRCA, we represented this in our model as three binary covariates, with a sample taking on a value of 1 in one of these covariates and a 0 in the two others.
C. <u>METABRIC: Comparison of Betas to TCGA-BRCA.</u> For each driver gene mutated at ≥5% frequency in both TCGA-BRCA and METABRIC cohorts, which includes *TP53* and *PIK3CA*, we computed the Spearman correlation between the fit coefficients for all target genes tested in both TCGA-BRCA and METABRIC (Fig. 3B, Fig. S5C).

### VII. Gene Set Enrichment Analysis

We compute enrichment in curated gene sets (Fig. 2B-D; Table S3) using a preranked, one-sided gene set enrichment test with the fgsea package’s *fgseaMultilevel* function^111^, which uses a multilevel splitting Monte Carlo approach and provides an enrichment score, normalized enrichment score, and *p*-value for each gene set. We use gene sets from the DoRothEA network^28^ (Fig. 2B), a curated set of *TP53* targets from Fischer et al.^29^ (Table 1, Fig. 2C) and Reactome^41^ (Fig. 2C), and from the CGC^33^ (Fig. 2D). The DoRothEA network was accessed via the dorothea R package using the *dorothea_hs* function. Only high confidence genes (levels A-C) were used, as has been done in previous work. Additional gene sets used for *TP53*-specific gene set enrichment analysis were downloaded directly from TRRUST^30^, hTFtarget^31^, and KEGG^32^ websites (Table S3).

We compute pathway-level gene set enrichment analysis (Fig. S1C; Data S2) using the package ReactomePA^112^ in R, specifically using the *gseGO, gseKEGG,* and *gseMKEGG* functions applied to the –*log*(*q*-values) multiplied by the directionality of the associated mutation coefficient (1 for coefficient > 0, –1 for coefficient < 0) produced by Dyscovr. In cases with more than five significant GO pathways, functionally similar GO pathways^34^ were consolidated using the Wang et al. method^113^; pathways with a similarity metric greater than 0.7 were merged, retaining the name of the more statistically significant pathway (Fig. S1C). Full sets of enriched GO pathways, without merging, can be found in Data S2.

### VIII. Identification of Genetic Interactions Using the Cancer Dependency Map (DepMap)^26^

We use a combination of CRISPRi gene dependency data, mutation data, and gene expression data from DepMap public v.23Q2 to narrow down putative candidates from phase I of Dyscovr to those that are potentially involved in a genetic interaction with the corresponding driver gene. For a given cancer driver gene *d,* we first limit *d*’s transcriptional hits to most confident hits, i.e. those that are statistically significant both pan-cancer (*q* < 0.2) and within at least one individual cancer type (*q* < 0.2). For each of these remaining targets *t* in set *T*, we use the following regression framework to relate the cell viability upon CRISPR knockout of *t* (*V*_*t*_) to the mutation status (*U*_*d*_) and expression (*E*_*d*_) of *d* across a set *K* of cancer types (*X*_*i*_) (see Methods VIII.A.d):

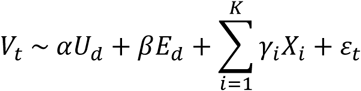

Overall, |*T |* regression models are fit for a given *d.* For both oncogenes and tumor suppressors, mutation status and expression provide complementary but non-equivalent measures of driver state, because activating or disruptive mutations can have downstream consequences that differ from those produced by increased or reduced expression. Thus, to uncover genetic interactions for driver gene *d*, we evaluate the significance of the combined effect of *U*_*d*_ and *E*_*d*_ on *V*_*t*_ using Empirical Brown’s method (i.e. the *empiricalBrownsMethod* function from the EmpiricalBrownsMethod R package with default parameters). This method is an adaptation of Fisher’s method^114^ that allows for term interdependency. With this approach, we obtain a single *p*-value for each potential genetic interaction considered. We obtain multiple hypothesis corrected *q*-values from these *p*-values using the *qvalue* function from the qvalue package in R with default parameters^88^.

For each driver gene *d*, we further classify which of its genetic interactions correspond to conditional genetic vulnerabilities with potential therapeutic relevance. In the case of an oncogene (*KRAS* and *PIK3CA*), we expect activating nonsynonymous mutation or high expression to result in cancer-promoting overactivity. Thus, promising co-targeting vulnerability candidates (i.e., cases where joint inhibition of the driver and the predicted interactor reduces cell viability) would display a positive relationship between activity of *d* and cell viability upon knockout of *t* (α > 0, β > 0) (Fig. 4A). Conversely, a promising acquired vulnerability candidate (i.e., cases where inhibition of the predicted interactor reduces cell viability in the context of a mutant driver) would display a negative relationship between the activity of *d* and cell viability upon knockout of *t* (α < 0, β < 0). For tumor suppressor genes (*TP53*), we search only for acquired vulnerabilities, as cancer-relevant nonsynonymous mutations in tumor suppressors impair their tumor-suppressor activity (and thus, further inhibiting an already impaired tumor suppressor in combination with the target gene would not represent a plausible therapeutic strategy). In this case, we expect a negative relationship between the mutation status of *d* and cell viability upon knockout of *t*, but a positive relationship between the expression of *d* and cell viability upon knockout of *t* (α < 0, β > 0). We define oncogenes and tumor suppressors using the standard Vogelstein definitions^27^. We report the set of targets *t* for which the combined *q*-value is less than 0.2 and the coefficients α and β from the pre-recombined analysis align with the above schema (Table S4). We note that for *TP53* and tumor suppressors more generally, α > 0, β < 0 corresponds to interactions in which greater tumor suppressor activity is associated with greater sensitivity to target loss (i.e. in which the presence of functional *TP53* increases dependency on the interacting gene); this case does not correspond to a plausible therapeutic strategy for *TP53*-mutant tumors. Combinations of α and β other than those listed above indicate discordant associations of driver mutation status and expression with sensitivity to target loss. These patterns may reflect biologically distinct consequences of driver mutation and altered driver expression on sensitivity to target loss, and are therefore not assigned to either conditional-vulnerability class.

A. <u>DepMap: Data Acquisition and Processing.</u> CRISPR dependency, mutation, gene expression, and cell line metadata for cancer cell lines were downloaded from the DepMap online portal (https://depmap.org/portal/). Models were run using the 693 cancer cell lines with all data types available and the set of 3 TCGA pan-cancer drivers (*TP53, PIK3CA,* and *KRAS*) that have nonsynonymous mutations in at least 15 cell lines across cell lines in the set of cancer types in which that driver is recurrently mutated (Table S1).
  a. *<u>CRISPR Dependency Data</u>.* Each target gene *t*’s dependency score upon genetic interference corresponds to the CRISPR dependency value *V*_*t*_. Dependency scores were processed using Chronos^115^. Positive dependency values indicate increased cell viability upon knockout, while negative dependency values indicate reduced cell viability upon knockout.
  b. *<u>Expression Data</u>. log2(*TPM + 1)-normalized RNA-seq files from DepMap were quantile normalized so that each of the cell lines has the same distribution of gene expression values using scikit learn’s *quantile.transform* function^92^, with an output distribution of ‘normal.’ Each driver gene *d*’s expression level corresponds to a quantile-normalized gene expression value *E*_*d*_.
  c. *<u>Mutation Data</u>.* The DepMap mutation file was subsetted to include only nonsynonymous mutations, selecting for ‘VariantInfo’ column annotations that include ‘MISSENSE’, ‘NONSENSE’, ‘NONSTOP’, and ‘SPLICE_SITE’. The mutation status of each driver gene *d* in each sample, *U*_*d*_, is 1 if the driver gene has a nonsynonymous mutation in that sample and 0 otherwise.
  d. *<u>Cancer Type</u>.* Consists of a binary covariate for each cancer type. For any given cell line, one of these covariates will take on a value of 1 and all others a value of 0, depending on which cancer type it is classified as in the cell line metadata supplement column ‘primary_disease’. Cell line data from a given cancer type is only considered if *d* is mutated in at least 5% of samples from that cancer type in the TCGA cohort (Table S1).
B. <u>Assessment of Enrichment of Gold-Standard Genetic Interactions.</u> To validate Dyscovr’s predicted genetic interactors, we sought experimentally derived sources of genetic interactions with *TP53*, *PIK3CA*, and *KRAS*. For each of *TP53*, *PIK3CA*, and *KRAS*, we retrieved all genes from BioGRID^45^ v5.0.253 denoted as being involved in a genetic interaction with the given driver, considering any gene that met at least one of the listed classification codes for genetic interactions given in https://wiki.thebiogrid.org/doku.php/experimental_systems. This yielded a total of 423 genes for *TP53* (228 overlapping with our set of genes tested for genetic interactions), 52 genes for *PIK3CA* (45 overlapping), and 1941 genes for *KRAS* (1748 overlapping). Though Dyscovr predicted genetic interactors display strong enrichment in these genes on their own (one-sided GSEA, *TP53* NES = 1.33, *p* = 2.57E-04; *PIK3CA* NES = 1.33, *p* = 1.6E-01; *KRAS* NES = 1.16, *p* = 7.09E-02), we further supplemented our gene set with genetic interactors from literature sources specific to our drivers of interest (Table S5). For our final gene set, we took the total unique set of genes supported by any of the above sources, yielding a total of 946 genes for TP53 (856 overlapping with our tested set), 213 genes for PIK3CA (195 overlapping), and 5166 genes for KRAS (4576 overlapping). All enrichment tests were performed using fgsea, as in Methods VII. We did not further restrict this reference set to test specifically for enrichment amongst Dyscovr’s predicted conditional genetic interactions. While BioGRID further categorizes genetic interactions into various types of positive and negative genetic interactions, these classifications do not capture the driver context in which the interaction was measured. The underlying studies include those designed to identify interactions between wild-type drivers and target genes (e.g. high-throughput CRISPR knockout or shRNA screens such as Diehl *et al.*^116^); those focused specifically on interactions with mutant copies of a given driver (e.g. Xie *et al.*^117^); those that test both wild-type and mutant contexts (e.g. Lü *et al*.^118^); and those focused on drug-dependent interactions (e.g. Liu et al.^119^). Consequently, the reported interactions could not be consistently classified as acquired or co-targeting vulnerabilities. We instead perform a separate validation using the SynLethDB v2.0 database^46^, which catalogues SL interactions in cancer, the most-studied form of conditional genetic vulnerability. For each driver gene (*TP53, PIK3CA*, and *KRAS*), we retrieved all SL partners listed in SynLethDB. Because SynLethDB does not consistently annotate the molecular context in which each interaction was observed (e.g., mutant versus wild-type driver background), we mapped SynLethDB entries to the Dyscovr interaction classes that most closely correspond to the experimental designs represented in the database. For *TP53*, we considered both predicted acquired vulnerabilities (dependencies associated with *TP53* loss) and predicted interactions in which higher *TP53* activity was associated with increased sensitivity to target loss, reflecting SL relationships that arise in *TP53*-wild-type contexts. For the oncogenes *PIK3CA* and *KRAS*, we restricted analysis to Dyscovr’s predicted co-targeting vulnerabilities, corresponding to interactions involving simultaneous perturbation of the driver and the partner gene. To evaluate whether Dyscovr recapitulates SL relationships catalogued in SynLethDB, we performed gene set enrichment analysis using the ranked lists of Dyscovr interaction scores for each driver. Enrichment was assessed separately for two SynLethDB-derived gene sets: (i) SL interactions supported by experimental evidence or literature text mining, and (ii) SL interactions predicted by computational methods curated in SynLethDB. Enrichment tests were performed using the fgsea package as described above (Methods VII), using one-sided tests to assess whether SynLethDB SL partners were enriched among higher-ranked Dyscovr predictions. Only genes present in the Dyscovr-tested gene universe for each driver were included in the enrichment analysis.
C. I<u>dentification of Overlap Between Predicted Genetic Interactors and Known Drug Targets.</u> To identify predicted genetic interactors with existing drugs that target them, we retrieved an exhaustive set of gene-drug links from Santos *et al.* Supplemental Data S2^74^. We restricted this table to links involving *Homo sapiens* and extracted any entries where our predictions (*q* < 0.2) had at least one listed drug (Table S6). Genes were denoted as inhibitors/ antagonists via manual curation from the DrugBank database^120^.
D. <u>Evaluation of Candidate Genetic Interactions via Correlation to Drug Sensitivity.</u> For putative genetic interaction target *t* of clinical interest (i.e. *KBTBD2*), we further used DepMap’s repository of drug sensitivity data to compute a Spearman’s correlation between the viability of the cell line upon CRISPR interference of *t* (*V*_*t*_) and the sensitivity to each of 4659 drugs. Associated *p*-values were corrected using the qvalue package in R with default parameters^88^. Significant correlation to sensitivity of drugs (*q* < 0.2) were used to infer potential mechanisms-of-action of *t*.

### IX. Survival Analysis for *PIK3CA* Mutation and *KBTBD2* Expression

To evaluate whether the expression level of putative target gene *KBTBD2* (*E*_*K*_) has an effect on patient survival (*S*), taking into consideration the binary nonsynonymous mutation status of cancer driver gene *PIK3CA* (*U*_*P*_), we used the following Cox proportional hazards model:

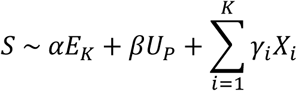

As in the main Dyscovr framework, we also have a number of additional covariates that correspond to various other clinical and molecular features that may have nontrivial effects on *S*, with the number *K* of such covariates dependent upon the characteristics of the set of samples being examined (i.e. pan-cancer or breast cancer, Methods IV.F). These covariates include the cancer type and subtype, age, sex, genotypic background, prior malignancies, prior treatment, nonsynonymous tumor mutational burden (TMB), tumor purity, and fraction of infiltrating immune cells. This model was fit using the *coxph* function in the survminer package in R version 0.4.9^121^ with default parameters. Hazard ratio estimates were computed as *e*^α^, where α is the fit coefficient for the *E*_*K*_ term (above).

A. <u>Data Acquisition and Representation</u>. Survival, mutation, and gene expression data were obtained for all 785 pan-cancer patients (see Methods IV.A and IV.B for details about mutation and gene expression data acquisition and preprocessing) from the National Cancer Institute’s GDC Data Portal with reported information about survival status and days until death or last follow-up (see Methods IX.A.a) and have high or low *KBTBD2* expression (see Methods IX.A.c).
  a. *<u>Survival Data.</u>* Survival data was extracted from the TCGA clinical supplement, including (a) binary survival status (alive or dead), and (b) time in days, defined as time until death if patient has died, or time until last follow-up if patient is alive. Survival *S* was represented as a *Surv* object from the survminer package for modeling.
  b. *<u>Mutation Data</u>.* The mutation status of *PIK3CA* in each patient, *U*_*P*_, is 1 if *PIK3CA* has a nonsynonymous mutation in that patients’ sample and 0 otherwise.
  c. *<u>Expression Data</u>*. The quantile-normalized expression matrix was further *z*-score normalized per patient column, such that the resulting matrix took on a mean (μ) of ∼0 and a standard deviation (σ) of ∼1. This formulation allowed us to select patients whose normalized expression value for *KBTBD2* is less than μ − σ (which we define as having “low” expression) or is greater than μ + σ (which we define as having “high” expression). Patients who do not fall into either of these categories are excluded from further analysis. In the Cox proportional hazards model, the expression status of *KBTBD2* in each patient, *E*_*K*_, is 1 if the patient has “high” *KBTBD2* expression and 0 if the patient has “low” *KBTBD2* expression, as previously defined.
B. *<u>Survival Visualization.</u>* Adjusted patient survival curves were visualized using the *ggadjustedcurves* function from the survminer package using the default ‘single’ (average for population) approach^121^. Briefly, this approach displays expected survival curves calculated based on the Cox model fit.

### X. Cell Lines

MCF7 (HTB-22) was purchased from American Type Culture Collection (ATCC) and maintained in DMEM supplemented with 10% Fetal Bovine Serum (FBS) and 1% penicillin and streptomycin. MCF7 cells possess an E545K hotspot *PIK3CA* mutation.

### XI. Immunoblotting

Cells were collected in ice cold PBS and lysed with RIPA lysis buffer (Pierce #89901) supplemented with Halt protease and phosphatase inhibitors (Pierce Chemical). Lysates were centrifuged at 20,000 × g for 5 minutes at 4°C. The supernatant was collected, and protein concentration was determined using the BCA kit (Pierce) per manufacturer’s instructions. Equal amounts of protein (20μg) in cell lysates were separated by SDS–PAGE, transferred to nitrocellulose membranes (GE healthcare), immunoblotted with specific primary and secondary antibodies and detected by chemiluminescence with the ECL detection reagents from Thermo Fisher or Millipore.

### XII. Antibodies

pAKT S473 (Cell Signaling Technology (CST) 4060), pS6 S235/S236 (CST 4858), p4EBP1 S65 (CST 9451), Cyclin D1 (CST 55506), p85 alpha (CST 4292), Beta Actin (CST 4967). Secondary Goat anti-Rabbit IgG (H+L) Secondary Antibody HRP (Thermo Fisher 65-6120).

### XIII. siRNA Knockdown

Cells were transfected for 48hr with Dharmacon SMARTpool nontargeting or siRNA designed against human KBTBD2. Transfection was aided by preincubation of siRNA with lipofectamine RNAiMAX (Thermo Fisher Scientific) and used according to the manufacturer’s instructions.

### XIV. mRNA extraction and RT-qPCR

mRNA was isolated using RNeasy kit (Qiagen) with Qiaschredder and eluted in 50uL. cDNA was synthesized using SuperScript First-Strand Synthesis System for RT-PCR kit (ThermoFisher #11904018). Synthesis was performed using random primers. Probes and primers were obtained from ThermoFisher as follows: KBTBD2 (Assay ID: Hs01556149_m1, FAM-MGB and GAPDH (Assay ID: Hs02786624_g1, VIC-MGB). qPCR was performed in 20uL reaction volumes with *KBTBD2* assay together with *GAPDH* and amplified using iTaq universal probes supermix (Bio-Rad Laboratories, #1725134). Relative quantification was performed using (2–ΔΔCt).

### XV. Quantification of Cell Growth and Viability

MCF7 cells were seeded into 24-well plates at 50,000 cells per well and transfected as indicated above. Cell growth was quantified using the sulforhodamine B assay. For each condition at least 3 replicates were measured. Growth at day 0 was subtracted from all Day 4 values and growth was normalized to Veh. treated samples.

## Resource Availability

Further information and requests for resources and reagents should be directed to and will be fulfilled by the lead contact, Mona Singh.

## Materials Availability

This study did not generate new unique reagents.

## Data and Code Availability

● This paper analyzes existing, publicly available data. These accession numbers for the datasets are listed in the key resources table.
● All original code has been deposited at GitHub and is publicly available as of the date of publication.
● Any additional information required to reanalyze the data reported in this paper is available from the lead contact upon request.

## Additional Resources

Dyscovr model results from TCGA patients are publicly searchable and downloadable at: dyscovr.princeton.edu.

