## Supplemental Materials for "Driver-associated transcriptional rewiring reveals conditional genetic vulnerabilities in cancer"

### Supplemental Information

Table S1: Vogelstein drivers mutated at ≥5% frequency per TCGA cancer type.

Table S2: *PIK3CA, KRAS,* and *IDH1* pan-cancer enrichments in effector TF targets.

Table S3: *TP53* pan-cancer enrichments in gold-standard target sets.

Table S4: Number of predicted genetic interactions with positive and/or negative driver covariates.

Table S5: Literature-curated genetic interaction gene sets.

Table S6: Predicted genetic interactors with FDA-approved drugs.

Table S7: Top ten predicted co-targeting vulnerabilities for *PIK3CA*.

Table S8: Number of putative target genes with significant pan-cancer associations to each regression term.

Table S9: Parameters for GDC Data Portal file downloads.

Table S10: Per-cancer TCGA subtype information.

Data S1: Genes predicted to be transcriptionally dysregulated by nonsynonymous mutations in *TP53*, *PIK3CA*, *KRAS*, and *IDH1*.

Data S2: GO and KEGG per-driver pathway enrichment.

Data S3: Variables removed due to multicollinearity, pan-cancer.

Data S4: Genes predicted to be transcriptionally dysregulated by driver mutations in individual cancer types.

Data S5: Predicted genetic interactors of *TP53*, *PIK3CA*, and *KRAS*.

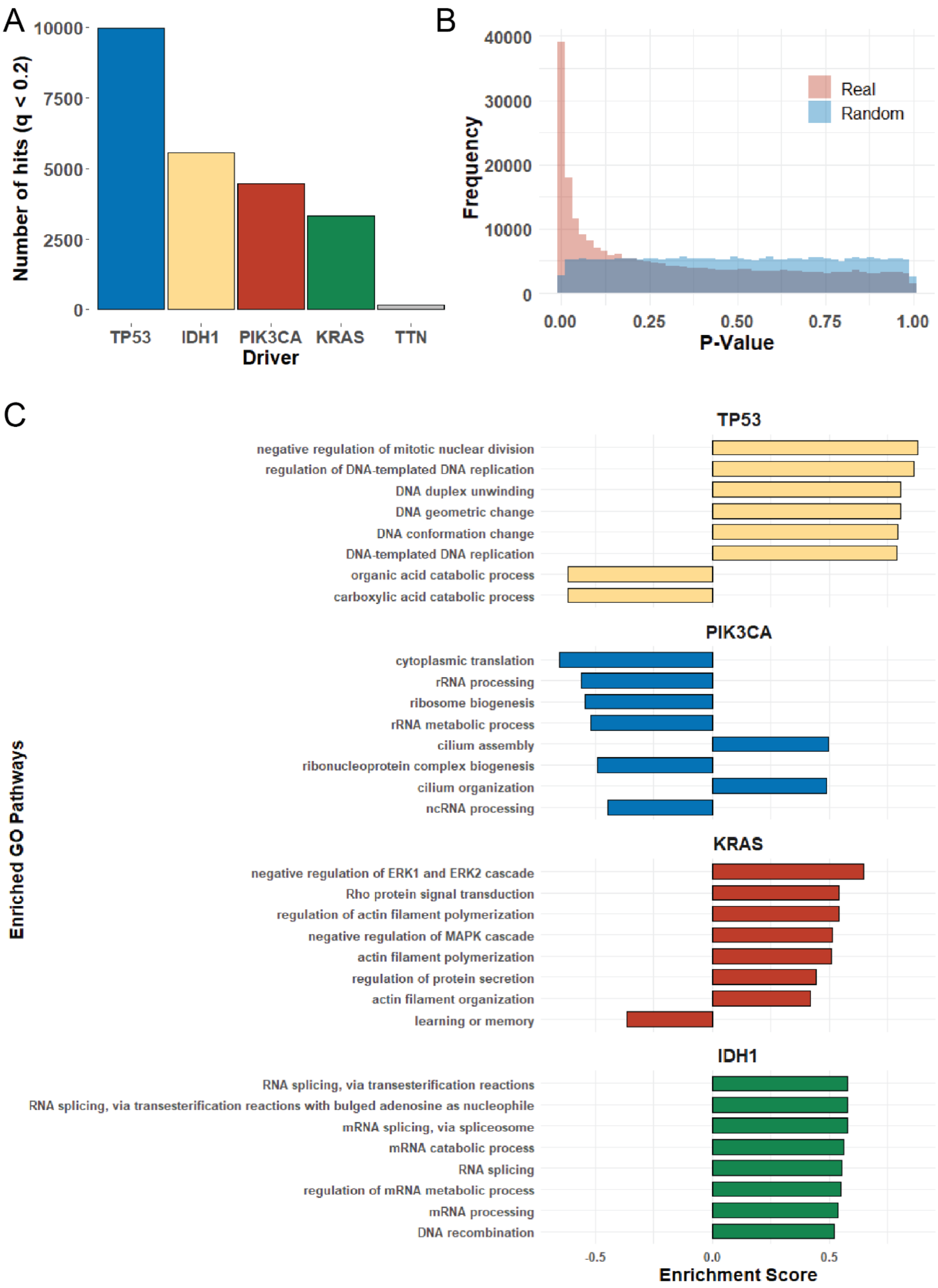


**Supplemental Figure 1. Pan-cancer quantitative validations and gene set enrichment analyses. A.** The number of genes predicted by the first phase of Dyscovr to be transcriptionally dysregulated in association with mutation in *TP53*, *PIK3CA*, *KRAS*, and *IDH1* in a pan-cancer setting ($q < 0.2$). Each hit represents a putative association between a nonsynonymous mutation in the driver gene and an expression change in a target gene.

Dyscovr identifies thousands of significant associations at this threshold for all recurrently mutated driver genes, whereas *TTN* (a recurrently mutated non-Vogelstein[1] or CGC[2] gene) has 160 hits at the same threshold. **B.** The distribution of *p*-values produced by Dyscovr for all driver mutation terms when applied pan-cancer across all candidate human gene targets (pink). Overlaid is the distribution of *p*-values obtained when expression values for each target gene are randomly permuted across samples prior to model fitting (blue, Methods IV.A). **C.** Bar charts of Gene Ontology (GO)[3] gene set enrichment analysis (GSEA) results for each driver gene using ReactomePA (Methods VII). Pathways were restricted to those enriched at $q < 0.05$ and ranked by -log10(*q*-value) multiplied by the direction of enrichment, with ties broken by descending leading-edge percentage. The top eight pathways for each driver are shown, with bar size corresponding to the enrichment score.

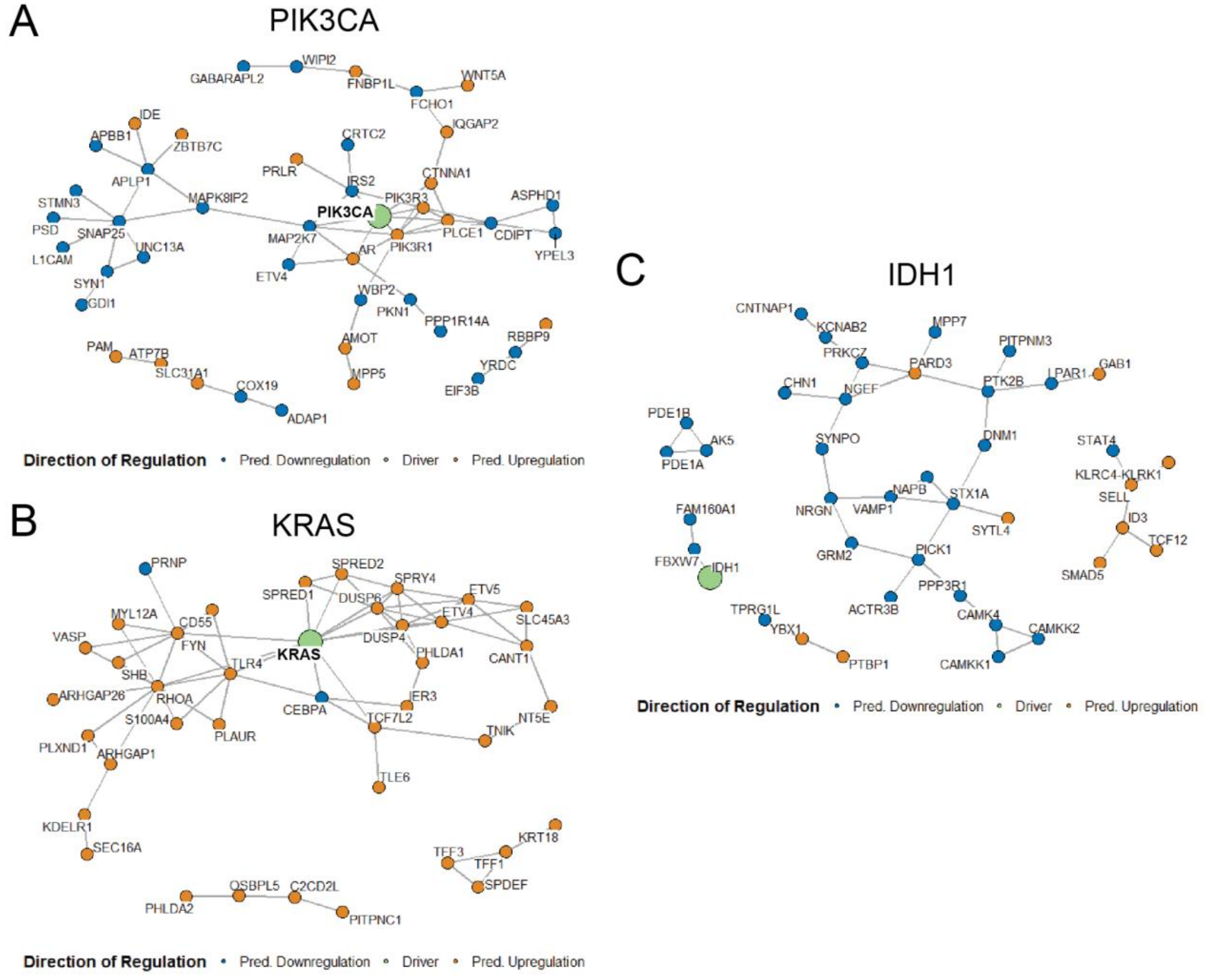


**Supplemental Figure 2. Overlay of pan-cancer hits from the first phase of Dyscovr ($q < 0.01$, Top 100) on the STRING functional network[4]. A-C.** Overlay of Dyscovr phase one results for *PIK3CA, KRAS,* and *IDH1* (respectively) on the STRING functional protein-protein interaction network. The corresponding driver gene is shown in green. The top 100 significant hits ($q < 0.01$) that are connected either to the driver or to another hit with STRING confidence >0.4 are displayed. Node color indicates the predicted direction of regulation (upregulation in orange, downregulation in blue). Disconnected components of size 2 or smaller are omitted from the visualization.

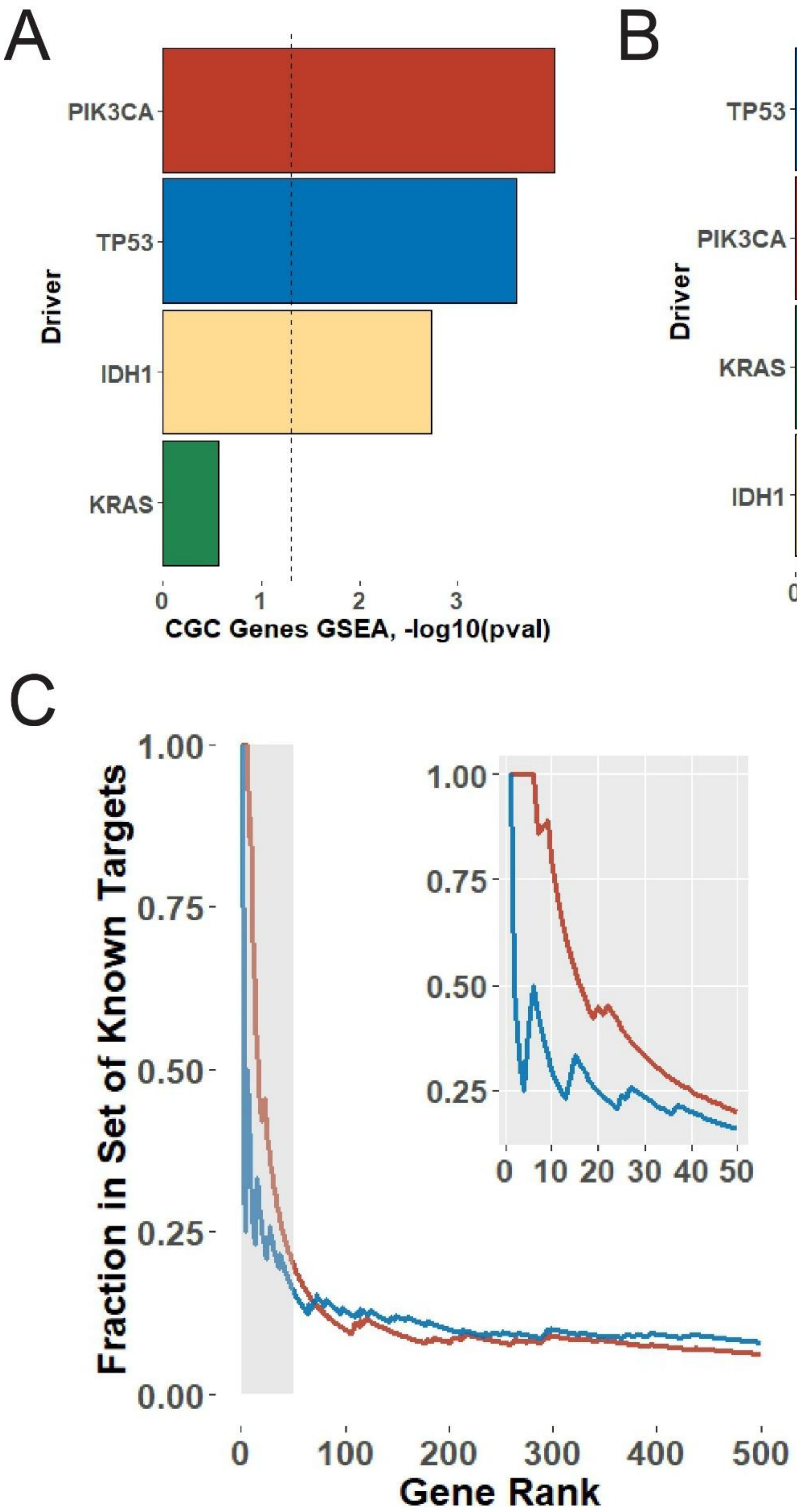


**Supplemental Figure 3. Evaluation of pan-cancer Dyscovr model using hotspot or nonsense mutation aggregation.** In the main analysis in our manuscript, we assumed that most mutations in *KRAS*, *PIK3CA,* and *IDH1* are oncogenic, as the majority of somatic mutations in these genes occur in reported 'hotspot' residues (in our TCGA cohort, 98% of *KRAS* mutations occur at G12, G13, and Q61[5]; 62% of *PIK3CA* mutations occur at E542, E545, H1047, and H1047[6]; and 83.6% of *IDH1* mutations occur at R132[7]). We also assume most mutations in *TP53* disrupt tumor suppressor function, as the majority of somatic variants in *TP53* are missense (75.0%) or nonsense (15.2%) and are thus expected to impair normal tumor suppressive activity, though some missense variants may also confer oncogenic properties[8]. In this figure, we restrict analysis to hotspot mutations in oncogenes (Methods IV.B.a) and nonsense or Clinical Interpretation of Variants in Cancer (CIVIC)[9]-supported missense mutations in *TP53*. **A.** Bar chart showing the enrichment (one-sided GSEA test) of known cancer genes from the CGC[2] in each driver gene's list of dysregulated targets ranked by Dyscovr phase one *q*-value. Bar height represents *-log10*(*p*-value), with dotted line denoting $p = 0.05$. **B.** Bar chart showing the enrichment of effector TF targets from DoRothEA[10] (confidence levels A-C), either *TP53*-specific targets for *TP53* (N = 248) or pooled targets across multiple, literature-supported effector TFs for *PIK3CA* (N = 605)*, KRAS* (N = 679)*, and IDH1* (N = 240) (Table S2)*.* Bar height represents *-log10*(*p*-value), with dotted line denoting $p = 0.05$*.* Enrichments correspond to the nonsense- or hotspot-specific version of Dyscovr described in panel A. **C.** The cumulative fraction of curated *TP53* targets (Fischer et al.[11], blue, one-sided GSEA $p = 2.79E-17$, NES = 1.74) and *TP53* signaling pathway members (Reactome[12] R-HSA-3700989, red, one-sided GSEA $p = 1.89E-25$, NES = 1.44) among an increasing fraction of *q-value* ranked *TP53* hits from a Dyscovr phase one model using only nonsense and CIVIC-supported missense mutations. Inlay (top right) shows the top 50 *TP53* hits.

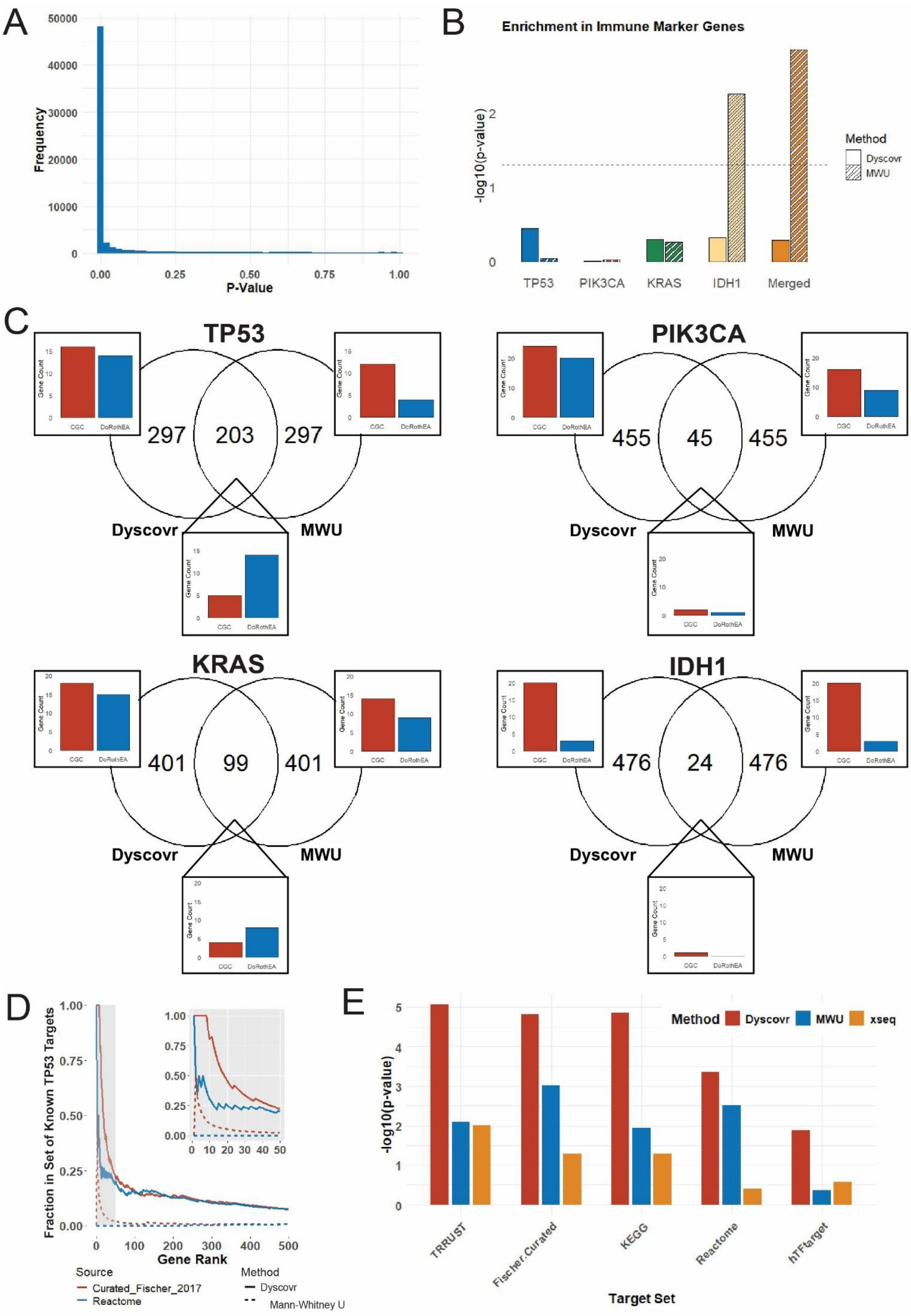

A
Frequency
P-Value
B
Enrichment in Immune Marker Genes
-log10(p-value)
Method
Dyscovr
MWU
TP53
PIK3CA
KRAS
IDH1
Merged
C
TP53
297
203
297
Dyscovr
MWU
PIK3CA
455
45
455
KRAS
401
99
401
IDH1
476
24
476
Gene Count
CGC
DoRothEA
D
Fraction in Set of Known TP53 Targets
Gene Rank
Source
Curated_Fischer_2017
Reactome
Method
Dyscovr
Mann-Whitney U
E
Method
Dyscovr
MWU
xseq
-log10(p-value)
TRRUST
Fischer.Curated
KEGG
Reactome
hTFtarget
Target Set

**Supplemental Figure 4. Comparison of Dyscovr to Mann-Whitney U (MWU) and xseq. A.** Histogram of *p*-values from MWU tests, consolidated across all four driver genes and all candidate dysregulated target genes. **B.** Bar plot showing the -log10(*p*-value) from a GSEA test for enrichment in immune cell marker genes[13] (N = 153) among candidate target genes ranked by *p*-value for each of *TP53*, *PIK3CA*, *KRAS*, and *IDH1*. Results are shown for Dyscovr phase one (solid bars) and MWU (striped bars) (Methods V.A). We also report enrichment significance for a merged pan-cancer ranking, where genes are ranked by the minimum *p*-value across the four drivers ("Merged"). MWU shows more significant enrichment in immune-related genes for *KRAS*, *IDH1*, and the merged condition. **C.** Venn diagrams showing overlap between the top 500 predicted dysregulated target genes, ranked by *p*-value, between Dyscovr phase one and MWU for each of *TP53*, *PIK3CA*, *KRAS*, and *IDH1.* Each segment contains an inset bar plot showing the number of genes in that segment that intersect either CGC genes (red) or DoRothEA TF target genes (blue). For DoRothEA, we use *TP53*-specific targets for *TP53* (N = 248) or pooled targets across multiple, literature-supported effector TFs for *PIK3CA* (N = 605)*, KRAS* (N = 679)*, and IDH1* (N = 240) (Table S2)*.* **D.** The cumulative fraction of curated *TP53* targets (Fischer et al.[11], blue) and *TP53* signaling pathway members (Reactome[12] R-HSA-3700989, red) among an increasing fraction of *p*-value ranked *TP53* hits from Dyscovr (solid lines) or MWU (dashed lines). Inlay (top right) shows the top 50 ranked *TP53* hits. **E.** For Dyscovr (red), MWU (blue), and xseq (orange), bars show the *-log10*(*p*-value) from one-sided GSEA tests for enrichment of known *TP53* target gene sets when restricted to genes present in xseq's provided interaction network. Gene sets include TRRUST[14] (N = 195), Fischer et al.[11] (N = 116), KEGG[15] hsa04115 (N = 69), Reactome[12] R-HSA-3700989 (N = 363), and hTFtarget[16] (N = 72).

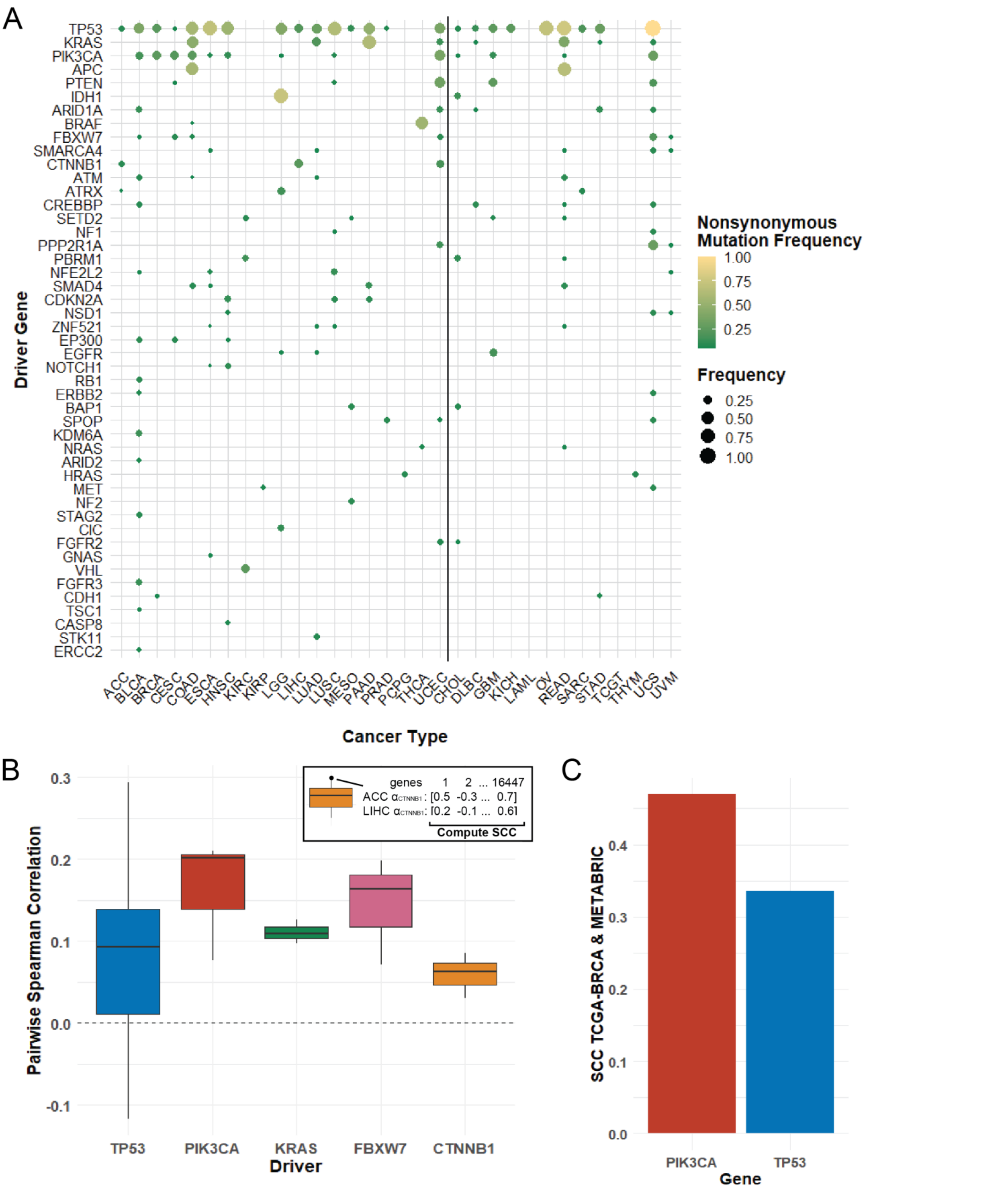


**Supplemental Figure 5. Driver mutation-target gene expression relationships across individual TCGA cancer types. A.** Cowplot showing the nonsynonymous mutation frequency for recurrently mutated (≥5%) driver genes, across TCGA cancer types. Each cancer driver gene annotated by Vogelstein[1] and mutated at ≥5%

frequency in at least one of the 19 TCGA cancer types with ≥75 samples meeting inclusion criteria (Methods II) is shown on the *y*-axis (left of vertical line). A dot indicates that the driver gene is mutated in ≥5% of the samples for the corresponding cancer type (*x*-axis). Dot size and color represent the nonsynonymous mutation frequency of that driver gene in that cancer type. Mutation frequencies of these drivers in cancer types with fewer than 75 samples are shown to the right of the vertical black line. **B.** Pairwise Spearman's rank correlations of estimated fit mutation coefficients for *TP53, PIK3CA*, *KRAS, CTNNB1,* and *FBXW7*, across all putative target genes. Correlations are computed across cancer types in which the given driver is frequently mutated (≥5% of samples) and has at least 15 significant dysregulated target genes ($q < 0.2$). The average Spearman's rank correlation for all drivers is greater than zero (dashed line), indicating that transcriptional effects of driver mutations show similarities across cancer types. **C.** Bar charts of the Spearman's rank correlation coefficient between the nonsynonymous mutation status coefficient for the indicated driver (*PIK3CA* in red, *TP53* in blue) across all putative target genes in TCGA-BRCA and METABRIC cohorts. The pairwise correlation coefficients are 0.47 for *PIK3CA* and 0.34 for *TP53* (both $p <$ 1E-200).

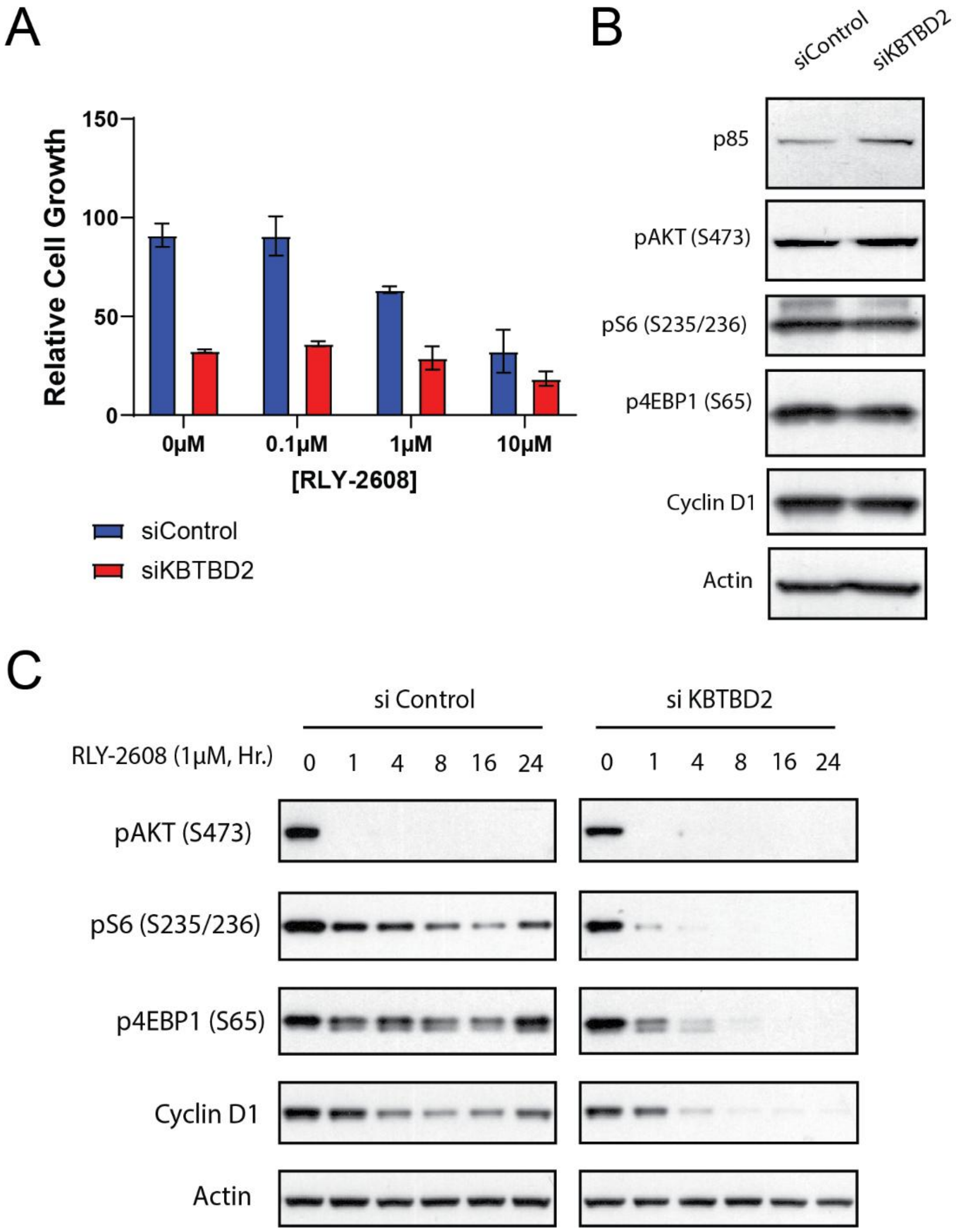


**Supplemental Figure 6. Characterization of *KBTBD2* in the context of mutant PI3K alpha inhibition. A.** Relative cell growth in response to RLY-2608 treatment in siControl and siKBTBD2 treated cells. MCF7 cells were transfected with siControl or siKBTBD2 for 48 hours followed by RLY-2608 treatment at indicated doses for an additional 2 days. Cell growth was quantified by SRB staining. Data are presented as mean ± SD. **B.** Knockdown of *KBTBD2* and western blot analysis of PI3K pathway components. MCF7 were transfected with control or siRNA targeting KBTBD2 for 48 hours. **C.** Western blot analysis of PI3K signal transduction following RLY-2608 treatment and *KBTBD2* knockdown. MCF7 cells were transfected with siControl or siKBTBD2 for 48 hours followed by treatment with 1 μM RLY-2608 for time *t* (hours).

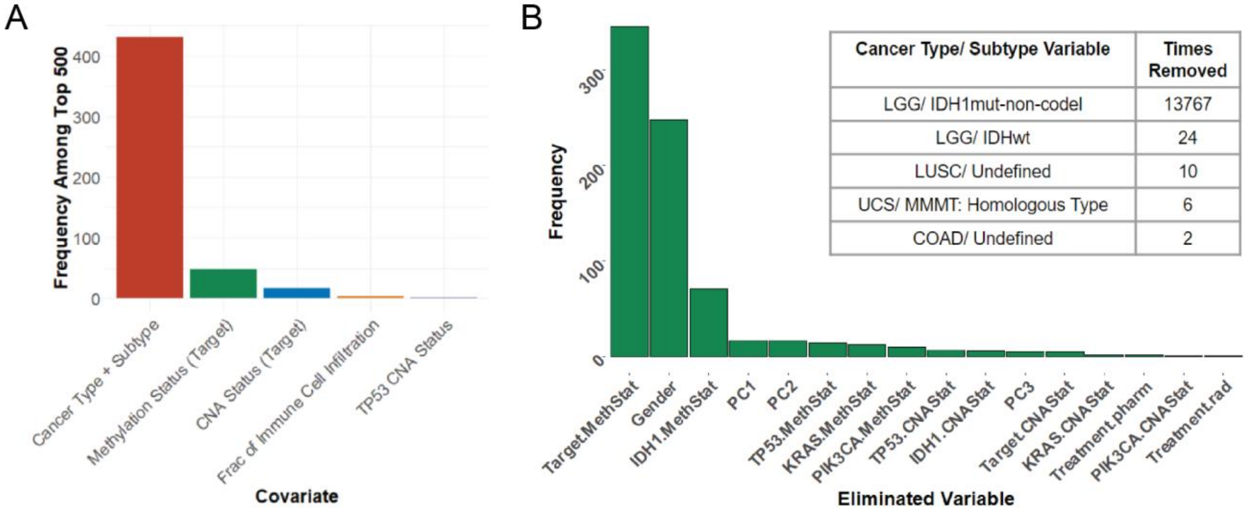


**Supplemental Figure 7. Pan-cancer Dyscovr covariate significance and covariates removed due to multicollinearity. A.** When the first phase of Dyscovr is applied in a pan-cancer setting across all putative gene targets, we aggregate all terms from these models together and rank them by *q*-value. Among the top 500 most significant terms from this ranking, we report the five most frequently represented covariate categories (when counting bucketed variables together, see Methods IV.F). The *x*-axis shows the covariate category and the *y*-axis shows its frequency among the top 500 terms. Cancer type and subtype variables are the most frequently represented, followed by the methylation status of the candidate target gene, the CNA status of the candidate target gene, immune cell infiltration fraction, and *TP53* CNA status. **B.** Bar plot (left) displays the number of times a given non-cancer type/ subtype covariate was removed from the regression framework due to multicollinearity when applying the first phase of Dyscovr pan-cancer across all genes (16,447 total regressions). Covariates were evaluated for collinearity using the variance inflation factor (VIF) and were removed if they had a VIF > 5. 'MethStat' denotes methylation status and 'CNAStat' denotes CNA status. PC1-3 correspond to germline principal components. The table (right) displays the number of times a given cancer type/ subtype covariate was removed due to collinearity. These variables were evaluated using Spearman correlation and were removed if their correlation with another covariate exceeded 0.7 with significance $p<1E-05$ (Methods III).

| **Cancer Type Abbrev.** | **Cancer Type Name** | **Vogelstein Driver Genes at ≥5% Frequency** | **Sample Size** |
|---|---|---|---|
| ACC | Adrenocortical carcinoma | *CTNNB1, TP53* | 76 |
| BLCA | Bladder Urothelial Carcinoma | *ARID1A, ARID2, ATM, CREBBP, EP300, ERBB2, ERCC2, FBXW7, FGFR3, KDM6A, NFE2L2, PIK3CA, RB1, STAG2, TP53, TSC1* | 349 |
| BRCA | Breast invasive carcinoma | *CDH1, PIK3CA, TP53* | 732 |
| CESC | Cervical squamous cell carcinoma and endocervical adenocarcinoma | *EP300, FBXW7, PIK3CA, PTEN, TP53* | 276 |
| CHOL | Cholangiocarcinoma | *NA* | 33 |
| COAD | Colon adenocarcinoma | *APC, FBXW7, KRAS, PIK3CA, SMAD4, TP53* | 236 |
| DLBC | Lymphoid Neoplasm Diffuse Large B-cell Lymphoma | *CARD11, MYD88, P2RY8, PIM1* | 36 |
| ESCA | Esophageal carcinoma | *GNAS, NFE2L2, PIK3CA, SMAD4, SMARCA4, TP53* | 158 |
| GBM | Glioblastoma multiforme | *EGFR, PIK3CA, PTEN, TP53* | 58 |
| HNSC | Head and Neck squamous cell carcinoma | *CASP8, CDKN2A, EP300, NOTCH1, NSD1, PIK3CA, TP53* | 465 |
| KICH | Kidney Chromophobe | *TP53* | 65 |
| KIRC | Kidney renal clear cell carcinoma | *PBRM1, SETD2, VHL* | 141 |
| KIRP | Kidney renal papillary cell carcinoma | *MET* | 224 |
| LAML | Acute Myeloid Leukemia | *NA* | 6 |
| LGG | Brain Lower Grade Glioma | *ATRX, CIC, EGFR, IDH1, PIK3CA, TP53* | 518 |
| LIHC | Liver hepatocellular carcinoma | *CTNNB1, TP53* | 359 |
| LUAD | Lung adenocarcinoma | *KRAS, STK11, TP53* | 361 |
| LUSC | Lung squamous cell carcinoma | *CDKN2A, NF1, NFE2L2, PIK3CA, PTEN, TP53, ZNF521* | 300 |
| MESO | Mesothelioma | *BAP1, NF2, TP53* | 81 |
| OV | Ovarian serous cystadenocarcinoma | *NA* | 7 |
| PAAD | Pancreatic adenocarcinoma | *CDKN2A, KRAS, SMAD4, TP53* | 170 |

| | | | |
|---|---|---|---|
| PCPG | Pheochromocytoma and Paraganglioma | *HRAS* | 167 |
| PRAD | Prostate adenocarcinoma | *SPOP, TP53* | 419 |
| READ | Rectum adenocarcinoma | *APC, KRAS, TP53* | 18 |
| SARC | Sarcoma | *TP53* | 58 |
| STAD | Stomach adenocarcinoma | *ARID1A, TP53* | 50 |
| TGCT | Testicular Germ Cell Tumors | *KIT* | 29 |
| THCA | Thyroid carcinoma | *BRAF, NRAS* | 396 |
| THYM | Thymoma | *NA* | 23 |
| UCEC | Uterine Corpus Endometrial Carcinoma | *ARID1A, CTNNB1, FBXW7, FGFR2, KRAS, PIK3CA, PPP2R1A, PTEN, SPOP, TP53* | 295 |
| UCS | Uterine Carcinosarcoma | *TP53* | 10 |
| UVM | Uveal Melanoma | *GNA11, GNAQ* | 17 |
| PC | Pan-Cancer | *TP53, PIK3CA, KRAS, IDH1* | 6135 |

**Supplemental Table 1. The Vogelstein et al. cancer driver genes**[1] **that are mutated in at least 5% of samples in each TCGA cancer type.** The driver genes mutated in at least 5% of pan-cancer samples are shown in the bottommost row.

| Driver Gene | Curated Effector TFs | # of Targets | One-Sided GSEA, *q*-value | ES | NES |
|---|---|---|---|---|---|
| *PIK3CA*[17] | *FOXO1/3/4/6* | 71 | 2.97E-01 | 0.50 | 1.15 |
| | *MYC* | 386 | 4.20E-01 | 0.42 | 1.03 |
| | *HIF1A* | 148 | 3.56E-01 | 0.45 | 1.08 |
| | *SREBF1/2* | 31 | **1.52E-03** | **0.69** | **1.54** |
| | *ATF4* | 16 | 5.87E-01 | 0.48 | 1.01 |
| | *NFR2/NFE2L2* | 11 | 8.03E-01 | 0.38 | 0.78 |
| *KRAS*[18–20] | *CREB1/3/5* | 117 | 5.46E-01 | 0.43 | 0.99 |
| | *MYC* | 386 | 5.42E-01 | 0.42 | 1.01 |
| | *ETS1/2* | 154 | **3.04E-06** | **0.61** | **1.43** |
| | *FOXO1* | 43 | **7.96E-03** | **0.62** | **1.37** |
| | *ELK1* | 31 | 5.42E-01 | 0.49 | 1.05 |
| | *ETV1* | 29 | **7.96E-03** | **0.67** | **1.45** |
| | *AP1 (absent from DoRothEA)* | N/A | N/A | N/A | N/A |
| *IDH1*[21] | *ATF3* | 50 | 1.56E-01 | 0.56 | 1.26 |
| | *MYCN* | 13 | 5.52E-01 | 0.47 | 0.97 |
| | *JUN/JUNB/JUND* | 176 | 5.52E-01 | 0.48 | 1.13 |
| | *NR2F2* | 18 | 2.96E-01 | 0.58 | 1.23 |
| | *SOX8 (no ABC confidence targets)* | N/A | N/A | N/A | N/A |
| | *HEY1 (no ABC confidence targets)* | N/A | N/A | N/A | N/A |

**Supplemental Table 2. Pan-cancer enrichment in transcriptional targets for effector TFs of *PIK3CA, KRAS*, and *IDH1*.** For each pan-cancer driver gene that is not itself a TF, we curated TFs shown in the literature to be effectors of that driver and tested for enrichment of each of those TFs' targets from DoRothEA[10]. The $q$-value from a multiple hypothesis corrected one-sided GSEA is shown for each effector TF ($q < 0.05$ in bold), along with the enrichment score (ES) and normalized enrichment score (NES).

| *TP53* Target List | # of Genes in Set | One-sided GSEA Test, *p*-value | ES | NES |
|---|---|---|---|---|
| ***Curated Targets and Databases*** | | | | |
| Curated, Fischer et al.[11] | 116 | 3.08E-16 | 0.81 | 1.69 |
| DoRothEA[10] (ABC) | 248 | 6.87E-17 | 0.72 | 1.54 |
| TRRUST[14] | 195 | 1.12E-15 | 0.73 | 1.56 |
| hTFtarget[16] | 72 | 1.41E-05 | 0.80 | 1.60 |
| ***Gene Pathways*** | | | | |
| KEGG Pathway[15] "P53 Signaling Pathway", hsa04115 | 69 | 2.35E-11 | 0.83 | 1.71 |
| Reactome Pathway[12] "Transcriptional Regulation by TP53", R-HSA-3700989 | 363 | 4.49E-14 | 0.66 | 1.42 |
| Reactome Pathway[12] "TP53 Regulates Metabolic Genes", R-HSA-5628897 | 83 | 9.29E-02 | 0.55 | 1.15 |
| ***Known Cancer Genes*** | | | | |
| CGC[2] Cancer Genes | 714 | 3.08E-03 | 0.51 | 1.12 |
| Vogelstein[1] Cancer Genes | 379 | 5.09E-02 | 0.51 | 1.10 |

**Supplemental Table 3. Enrichment of *TP53* pan-cancer dysregulated target genes from phase one of Dyscovr in curated gene sets, TF-target databases, and driver-specific genetic pathways.** For each gene set, we report the number of genes in each set, the enrichment significance, enrichment score (ES) and normalized enrichment score (NES) from a one-sided gene set enrichment test (Methods VII).

| Driver Gene | Category | Number of Genes ($q$ < 0.2) |
|---|---|---|
| TP53 | $\alpha > 0, \beta > 0$ | 345 |
| | $\alpha > 0, \beta < 0$ | 647 |
| | **$\alpha < 0, \beta > 0$ (acquired)** | **591** |
| | $\alpha < 0, \beta < 0$ | 436 |
| PIK3CA | **$\alpha > 0, \beta > 0$ (co-targeting)** | **33** |
| | $\alpha > 0, \beta < 0$ | 30 |
| | $\alpha < 0, \beta > 0$ | 68 |
| | **$\alpha < 0, \beta < 0$ (acquired)** | **43** |
| KRAS | **$\alpha > 0, \beta > 0$ (co-targeting)** | **70** |
| | $\alpha > 0, \beta < 0$ | 102 |
| | $\alpha < 0, \beta > 0$ | 65 |
| | **$\alpha < 0, \beta < 0$ (acquired)** | **60** |

**Supplemental Table 4. Number of putative pan-cancer genetic interaction pairings.** Reports the number of significant candidate genetic interactions (Methods VIII) categorized by the direction of association between the driver mutation status and cell viability following CRISPRi knockdown of the target gene ($\alpha$), and between driver expression and cell viability following CRISPRi knockdown of the target gene ($\beta$). The four categories correspond to ($\alpha > 0, \beta > 0$); ($\alpha > 0, \beta < 0$); ($\alpha < 0, \beta > 0$); and ($\alpha < 0, \beta < 0$). Categories corresponding to putative therapeutically relevant conditional genetic vulnerabilities for the given driver gene are highlighted in bold (Methods VII).

| Driver | Publication First Author (Year) | Number of Genes |
|---|---|---|
| *TP53* | Xie *et al.* (2012)[22] | 103 |
| | Imai *et al.* (2013)[23] | 50 |
| | Diehl *et al.* (2021)[24] | 39 |
| | Feng *et al.* (2022)[25] | 2034 |
| | Lü *et al.* (2024)[26] | 269 |
| *PIK3CA* | Diehl *et al.* (2021)[24] | 168 |
| | Naara *et al.* (2025)[27] | 32 |
| *KRAS* | Luo *et al.* (2009)[28] | 1306 |
| | Stekel *et al.* (2012)[29] | 77 |
| | Vizeacoumar *et al.* (2013)[30] | 386 |
| | Chang *et al.* (2016)[31] | 187 |
| | Martin *et al.* (2017)[32] | 2477 |

**Supplemental Table 5. Literature-curated genetic interaction gene sets.** Reports the number of genetic interactors reported in each of the following publications for each of *TP53, PIK3CA*, and *KRAS*.

| Driver Gene | Acquired or Co-targeting | Putative GI | Associated Drugs[33] |
|---|---|---|---|
| *TP53* | Acquired | *CACNA1D* | Amlodipine; Bepridil; Clevidipine; Diltiazem; Dronedarone; Felodipine; Isradipine; Nicardipine; Nifedipine; Nimodipine; Nisoldipine; Verapamil |
| | Acquired | *CFTR* | Crofelemer; Ivacaftor; Lumacaftor |
| | Acquired | *CHRNB1* | Atracurium; Cisatracurium; Decamethonium; Doxacurium; Gallamine; Metocurine; Mivacurium; Pancuronium; Pipecuronium; Rapacuronium; Rocuronium; Suxamethonium; Tubocurarine; Vecuronium |
| | Acquired | *DHFR* | Methotrexate; Pemetrexed; Pralatrexate |
| | Acquired | *EPHB4* | Vandetanib |
| | Acquired | *GHR* | Pegvisomant; Somatrem; Somatropin |
| | Acquired | *GRIA2* | Perampanel; Topiramate |
| | Acquired | *HDAC9* | Belinostat; Panobinostat; Romidepsin |
| | Acquired | *IMPA1* | Lithium carbonate; Lithium citrate |
| | Acquired | *KCNC1* | Dalfampridine; Guanidine |
| | Acquired | *KCNG3* | Dalfampridine; Guanidine |
| | Acquired | *KDR* | Axitinib; Cabozantinib; Lenvatinib; Nintedanib; Pazopanib; Ponatinib; Ramucirumab; Regorafenib; Sorafenib; Sunitinib; Vandetanib |
| | Acquired | *LAMA5* | Ocriplasmin |
| | Acquired | *PPARG* | Balsalazide; Mesalamine; Olsalazine; Pioglitazone; Rosiglitazone; Troglitazone |
| | Acquired | *PTH1R* | Parathyroid hormone; Teriparatide; Teriparatide acetate |
| | Acquired | *PTK6* | Vandetanib |
| | Acquired | *RYR3* | Dantrolene |
| | Acquired | *TUBB* | Brentuximab vedotin; Cabazitaxel; Colchicine; Docetaxel; Eribulin; Ixabepilone; Paclitaxel; Trastuzumab emtansine; Vinblastine; Vincristine; Vinorelbine base |
| *PIK3CA* | Co-targeting | *AR* | Bicalutamide; Danazol; Dromostanolone propionate; Enzalutamide; Ethylestrenol; Fluoxymesterone; Flutamide; Methyltestosterone; Nandrolone decanoate; Nandrolone phenpropionate; Nilutamide; Oxandrolone; Oxymetholone; Stanozolol; Testosterone; Testosterone cypionate; Testosterone enanthate; Testosterone propionate; Testosterone undecanoate |
| | Co-targeting | *IGF1R* | Mecasermin; Mecasermin rinfabate |
| *KRAS* | Co-targeting | *IFNGR2* | Interferon Gamma-1B |

| | | | |
|---|---|---|---|
| | Acquired | *ACE* | Benazepril; Captopril; Enalapril; Enalaprilat; Fosinopril; Lisinopril; Moexipril; Perindopril; Quinapril; Ramipril; Spirapril; Trandolapril |
| | Acquired | *CA9* | Dichlorphenamide; Ethoxzolamide |
| | Acquired | *CHRNA2* | Carbamazepine; Dextromethorphan |
| | Acquired | *EPHB4* | Vandetanib |

**Supplemental Table 6. FDA-approved drug targets among predicted Dyscovr genetic interactors for *TP53*, *PIK3CA*, and *KRAS*.** Predicted genetic interactors identified in phase two of Dyscovr ($q < 0.2$) that correspond to established drug targets are listed, along with associated US FDA-approved drugs[33]. The table includes both activators and inhibitors of the indicated genes; drugs with known inhibitory mechanisms-of-action (including antagonists) are highlighted in orange.

| **Predicted co-targeting conditional genetic vulnerability** | ***q*-value** |
|---|---|
| *CNPPD1* | 5.31E-03 |
| *TM4SF1*[34] | 1.28E-02 |
| *SUFU*[35] | 1.53E-02 |
| *IRS2*[36] | 2.04E-02 |
| *SLC7A2*[37] | 2.27E-02 |
| *IGF1R*[38] | 2.90E-02 |
| *HMGN4* | 3.85E-02 |
| *KBTBD2* | 6.10E-02 |
| *AKR1D1* | 6.50E-02 |
| *TCF7L2*[39] | 6.50E-02 |

**Table S7. Top predicted co-targeting genetic vulnerabilities for *PIK3CA*.** The ten most statistically significant predicted co-targeting genetic vulnerabilities for the cancer driver gene *PIK3CA* (ranked by *q*-value) identified by phase two of Dyscovr. Genes with previously reported relationships to *PIK3CA* or PI3K-AKT signaling are highlighted in orange.

| Regression Term | Number of Hits ($q < 0.01$) | Number of Hits ($q < 0.2$) |
|---|---|---|
| Cancer Type/ Subtype | 15,749 | 16,442 |
| Target Gene CNA Status | 13,577 | 16,441 |
| Target Gene Methylation Status | 13,095 | 16,098 |
| Total Immune Cell Infiltration Fraction | 11,782 | 16,206 |
| Tumor Purity | 9,455 | 15,471 |
| PIK3CA Methylation Status | 6,145 | 13,011 |
| KRAS Methylation Status | 5,928 | 12,585 |
| PIK3CA CNA Status | 5,678 | 12,452 |
| Germline PC1 | 5,595 | 11,969 |
| TP53 Mutation Status | 4,094 | 10,730 |
| TP53 CNA Status | 3,598 | 9,662 |
| KRAS CNA Status | 2,504 | 8,137 |
| IDH1 Methylation Status | 2,309 | 7,969 |
| TP53 Methylation Status | 2,064 | 7,765 |
| IDH1 CNA Status | 2,035 | 7,305 |
| Patient Age | 1,835 | 8,083 |
| Germline PC2 | 857 | 5,219 |
| IDH1 Mutation Status | 782 | 5,513 |
| KRAS Mutation Status | 400 | 3,317 |
| PIK3CA Mutation Status | 392 | 4,413 |
| Patient Sex | 363 | 4,303 |
| Target Gene Mutation Status | 42 | 201 |
| Germline PC3 | 2 | 17 |
| Tumor Mutational Burden (TMB) | 0 | 0 |
| Patient Treatment Status (Radiation) | 0 | 1 |
| Patient Prior Malignancy | 0 | 31 |
| Patient Treatment Status (Pharmaceutical) | 0 | 0 |

**Supplemental Table 8. Number of putative target genes significantly associated with each regression covariate in the pan-cancer Dyscovr phase one model.** At each indicated significance threshold, the table reports the number of genes whose expression is significantly associated with each covariate included in the first phase of the Dyscovr regression framework. For categorical variables (Methods IV.F), a reported hit indicates that the gene is significantly associated with at least one corresponding binary dummy covariate.

| Data Category | Data Type | Experimental Strategy | Workflow Type | Data Format | Platform |
|---|---|---|---|---|---|
| Simple nucleotide variation | Aggregated somatic mutation | WXS | MuSE Variant Aggregation and Masking | maf | N/A |
| Copy number variation | Gene level copy number | Genotyping array | ASCAT2 | txt | affymetrix snp 6.0 |
| Transcriptome profiling | Gene expression quantification | RNA-seq | HTSeq - Counts | txt | N/A |
| DNA methylation | Methylation Beta value | Methylation Array | Liftover | txt | illumina human methylation 450 |
| Biospecimen | Biospecimen supplement | N/A | N/A | bcr xml | N/A |
| Clinical | Clinical supplement | N/A | N/A | tsv | N/A |

**Supplemental Table 9. Parameters for GDC Data Portal file downloads.** Each column represents a facet of the data that can be selected on the GDC Data Portal website when downloading files. For each Data Category (column 1), specific data facet selections used in these analyses are provided for reproducibility.

| Cancer Type Abbrev. | Cancer Type Name | Column name in TCGAbiolinks clinical supplement | Subtypes |
|---|---|---|---|
| ACC | Adrenocortical carcinoma | COC | COC1, COC2, COC3 |
| BLCA | Bladder Urothelial Carcinoma | mRNA cluster | Luminal_infiltrated, Luminal_papillary, Luminal, Basal_squamous, Neuronal, ND |
| BRCA | Breast invasive carcinoma | BRCA_Subtype_PAM50 | LumA, LumB, Her2, Basal, Normal |
| CESC | Cervical squamous cell carcinoma and endocervical adenocarcinoma | SAMP:CIMP_call | CIMP-low, CIMP-intermediate, CIMP-high |
| COAD | Colon adenocarcinoma | MSI_status | MSI-H, MSI-L, MSS, Not Evaluable |
| DLBC | Lymphoid Neoplasm Diffuse Large B-cell Lymphoma | N/A | N/A |
| ESCA | Esophageal carcinoma | MSI status | MSI-H, MSI-L, MSS, NA |
| GBM | Glioblastoma multiforme | Original.Subtype | Classical, G-CIMP, IDHmut-codel, IDHmut-non-codel, IDHwt, Mesenchymal, Neural, Proneural |
| HNSC | Head and Neck squamous cell carcinoma | RNA | Atypical, Basal, Classical, Mesenchymal |
| KICH | Kidney Chromophobe | Histological.Subtype | Kidney Chromophobe |
| KIRC | Kidney renal clear cell carcinoma | N/A | N/A |
| KIRP | Kidney renal papillary cell carcinoma | tumor_type.KIRP.path. | Type 1 Papillary RCC, Type 2 Papillary RCC, Unclassified Papillary RCC |
| LGG | Brain Lower Grade Glioma | Original.Subtype | Classical, G-CIMP, IDHmut-codel, IDHmut-non-codel, IDHwt, Mesenchymal, Neural, Proneural |
| LIHC | Liver hepatocellular carcinoma | iCluster clusters (k=3, Ronglai Shen) | iCluster:1, iCluster:2, iCluster:3 |
| LUAD | Lung adenocarcinoma | iCluster.Group | 1,2,3,4,5,6 |
| LUSC | Lung squamous cell carcinoma | Expression.Subtype | basal, classical, primitive, secretory |
| MESO | Mesothelioma | N/A | N/A |
| PAAD | Pancreatic adenocarcinoma | Histological type by RHH | Ductal adenocarcinoma, Adenosquamous, Other, Colloid (mucinous noncystic) |
| PCPG | Pheochromocytoma and | mRNA Subtype Clusters | Kinase signaling, Wnt-altered, Pseudohypoxia, Cortical |

| | | | |
|---|---|---|---|
| | Paraganglioma | | admixture, NA |
| PRAD | Prostate adenocarcinoma | Subtype | 1-ERG, 2-ETV1, 3-ETV1, 4-FLI1, 5-SPOP, 6-FOXA1, 7-IDH1 8-other |
| READ | Rectum adenocarcinoma | MSI_subtypes | N/A |
| THCA | Thyroid carcinoma | mRNA_Cluster_number | 1,2,3,4,5,NA |
| THYM | Thymoma | N/A | N/A |
| UCEC | Uterine Corpus Endometrial Carcinoma | msi | Indeterminant, MSI-H, MSI-L, MSS |
| UCS | Uterine Carcinosarcoma | histologic subtype | Uterine Carcinosarcoma/ MMMT: Heterologous Type, Uterine Carcinosarcoma/ MMMT: Homologous Type, Uterine Carcinosarcoma/ Malignant Mixed Mullerian Tumor (MMMT): NOS |
| UVM | Uveal Melanoma | mRNA Cluster No. | 1,2,3,4 |

**Supplemental Table 10. Per-cancer TCGA subtype information.** For each TCGA cancer type, displays the TCGAbiolinks clinical supplement column name that contains the cancer subtype information used in Dyscovr (column 2) and the range of subtype values found in that column (column 3). In all cases, molecular subtypes were used, if available. If not, histological subtypes were preferentially used, followed by expression-based subtypes, such as from mRNA clusters.
